# Making Course Structure Visible in a Multi-Instructor Graduate Genomics Course: A Course-Level Evaluation of Standardized Learning Supports

**DOI:** 10.64898/2026.05.06.723173

**Authors:** Marie Saitou, Celian Diblasi

## Abstract

Graduate-level genomics courses require students to integrate dense material across subfields, concepts and methods. In modular, multi-instructor courses, students may struggle because the coherence between lectures can be difficult to navigate, while the course structure may be visible to instructors. We evaluated a 2025 navigation redesign of BIO322, a graduate genomics course at the Norwegian University of Life Sciences, while preserving course content, multi-instructor teaching, modular organization and assessment framework. The redesign includes introducing a standardized self-learning guide, expanded syllabus, enriched online quiz feedback, and added support for a final group research proposal. Using anonymized course evaluation scores from 2021–2025 and aggregated learning management system access data from 2023–2025, we examined student experience and resource use. In 2025, five of six course evaluation items reached their highest observed BIO322 scores, while one, lecture-specific score remained within the previous range. The consolidated self-learning guide was accessed by nearly all students, whereas access to optional readings declined across the course sequence, despite comparatively stable page views per accessing student. These course-level findings are consistent with improved perceived navigability following the introduction of standardized learning support. However, some students continued to report difficulty identifying priorities and connections among course components, indicating that challenges in perceived course coherence remained for part of the cohort despite the redesign.

**Practitioner Points:**

- Making course structure explicit may improve students’ perceived navigability in multi-instructor graduate genomics courses.
- A centralized self-learning guide can broaden access to preparatory guidance without changing core course content or assessment.
- Optional learning supports may be used unevenly, so resource availability should not be assumed to translate into uniform resource access.

## Introduction

Graduate-level genomics courses require students to engage with conceptually dense and rapidly changing material that is rarely reducible to a stable textbook sequence [1]. Students must learn concepts related to genome structure, evolutionary processes, population-level variation, gene regulation, and functional interpretation, while also encountering contemporary computational and statistical approaches used to analyze genomic data, as exemplified in several recent review articles [2–5]. Previous reports on genomics education have emphasized the need for course designs that support both conceptual understanding and resource access with research practices in genomics [6,7]. In this context, students must not only learn individual topics, but also understand how concepts from different genomics subfields connect to one another.

These demands can become particularly challenging in multi-instructor genomics courses, where different sessions are taught by instructors with distinct disciplinary expertise. Such teaching models can be pedagogically valuable because they expose students to up-to-date research perspectives and methodological diversity [8]. However, prior work on multi-instructor and team-taught courses indicates that students may experience such courses as fragmented when coherence, continuity, and expectations are not made sufficiently explicit [9,10]. Thus, the educational value of disciplinary breadth depends partly on whether students can perceive the course as a coherent learning trajectory rather than a sequence of separate expert-led sessions [11].

In advanced genomics courses, this issue is especially relevant because disciplinary diversity and instructional variation overlap with heterogeneous student backgrounds [12]. Students may enter the course with different prior exposure to genome biology, evolutionary reasoning, statistical interpretation, and data-driven analysis. As a result, the same lecture, reading, or assignment may pose different challenges for different students [13]. Even when a syllabus, learning objectives, and materials are provided, students may experience the course as difficult to navigate if expectations are distributed across documents, presented in inconsistent formats, or insufficiently connected to weekly activities [14–16]. From the perspective of cognitive load theory, such fragmented or poorly signposted information can increase extraneous cognitive load by requiring students to infer structure from dispersed cues rather than focusing on the underlying concepts [17].

Previous work in life sciences education has shown that structured course design, preparatory scaffolding, and low-stakes formative assessment can support student engagement and learning [18–20]. These approaches may reduce extraneous cognitive load, clarify expectations, and help students monitor their own understanding [17,21,22]. In multi-instructor genomics courses, such supports may also function as navigational tools, helping students identify what is central, what is supplementary, and how individual sessions relate to the broader course structure.

However, providing learning supports does not guarantee that students will recognize, prioritize, or use them as instructors intend. Students differ in prior knowledge, study strategies, time constraints, confidence, and ability to identify which resources are most relevant [23,24]. Student engagement with available supports should therefore be treated as an empirical outcome rather than assumed as a direct consequence of course design. This distinction is particularly important when learning supports are optional or preparatory, because their educational value depends not only on their design but also on whether students access and use them.

BIO322, a graduate-level genomics course at the Norwegian University of Life Sciences, provides a case in which these issues became visible through repeated course evaluations. The course serves approximately 70–80 master’s students from biology-related programmes and is taught by multiple instructors with different disciplinary expertise. It covers major areas of contemporary genomics, including genetic differences between species, genetic diversity among individuals within species, and gene function or regulatory differences within individuals. Because students enter the course with heterogeneous prior exposure to genome biology, statistical reasoning, and computational analysis, the course requires students to integrate concepts across several genomics subfields.

BIO322 underwent a major pedagogical restructuring in 2022, following the departure of the previous course responsible. The course was changed from a predominantly lecture-based format led by many instructors into a combination of lectures, hands-on group exercises, and module-level quizzes. The course structure was reorganized into three modules: genetic differences between species, genetic diversity among individuals within species, and gene function differences within individuals. To reinforce learning, quizzes were introduced at the end of each module, and hands-on genomic analysis exercises were developed with accompanying problem sets and solutions made available online. These resources were intended to support preparation, revision, and continuity across the course.

Despite this restructuring, course evaluation results varied across subsequent iterations. Student feedback repeatedly pointed to difficulties in preparing for individual sessions, prioritizing learning materials, understanding how topics connected across modules, and recognizing how lectures related to quizzes and assignments. These concerns occurred despite the presence of a formal syllabus, stated learning objectives, and established assessment formats. This indicated not an absence of structure, but a mismatch between instructor-intended structure and student-perceived navigability. This distinction aligns with work on constructive alignment and student perceptions of instructional materials, which suggests that alignment intended by instructors may not be experienced as alignment by students unless expectations and learning pathways are made accessible [24,25].

In response, the 2025 course iteration introduced a standardized self-learning guide for all sessions (**Supplementary File 1**). This guide replaced heterogeneous instructor-specific preparatory materials with a consistent format that identified lecture topics, key vocabulary, background concepts, guiding questions, and optional resources. The purpose was not to change the core content, modular organization, or assessment framework of the course. Instead, the intervention aimed to make the existing course structure more visible and easier to navigate. The guide was designed as preparatory scaffolding: a tool to help students identify what to attend to before and during each session, while preserving the disciplinary diversity of the multi-instructor format. It also served a bridging function across instructor-led sessions, similar to approaches in which a primary teacher or coordinating structure helps students perceive coherence in team-taught courses [26].

In parallel, administrative information was consolidated into a single expanded syllabus (**Supplementary File 2**), and online quizzes were revised to align more explicitly with module-level learning objectives. Mandatory quizzes were updated to include brief explanations for correct responses, thereby strengthening their formative function. Additional optional quizzes were introduced to support early-stage knowledge checking and preparation for the final group-based research design assignment. Because open-ended assignments can remain difficult for students who are still developing disciplinary expertise, these supports were intended to provide concrete entry points into the task [27,28]. These changes were intended to clarify expectations, support self-paced revision, and reduce navigation-related effort without altering the pass/fail assessment framework.

The present study examines how students engaged with course resources following this redesign, focusing on variation in aggregate patterns of student resource access. We use learning management system data and institutional course evaluation scores to characterize patterns of resource access in an authentic graduate genomics course setting. Because no consent responses were received for the use of individual-level learning management system or assessment data, the analysis was restricted to anonymized institutional evaluation results and aggregated course-level resource-access summaries.

Specifically, this study addresses three research questions. First, how did students’ perceived course navigability change after the introduction of standardized learning support? Second, how did access to a centralized course-wide guide compare with access to session-linked preparatory materials used in previous course iterations? Third, how did students access different categories of course resources after the redesign? By examining course-level evaluation patterns and aggregate behavioral resource access data, this study provides a course-level evaluation of a standardized learning-support redesign, focusing on perceived navigability and resource access.

## Materials and Methods

### Course Context and 2025 Redesign

BIO322 is a graduate-level genomics course offered at the Norwegian University of Life Sciences. The course serves approximately 70–80 master’s students from biology-related programmes and is taught by multiple instructors with different disciplinary expertise. The course is organized into three modules: genetic differences between species, genetic diversity among individuals within species, and gene function differences within individuals. The course includes weekly lectures, hands-on group exercises, module-level quizzes, and a final group-based research design assignment.

The 2025 intervention aimed to improve the navigability of the existing course structure without substantially changing the core content, modular organization, multi-instructor format, or pass/fail assessment framework. Administrative information was consolidated into an expanded syllabus, and academic preparatory information was consolidated into a standardized self-learning guide covering all teaching sessions. The guide identified lecture topics, key vocabulary, background concepts, guiding questions, preparatory cues, and optional resources in a consistent format across sessions. It was released before the start of the course (**Table 1**).

**Table 1.**
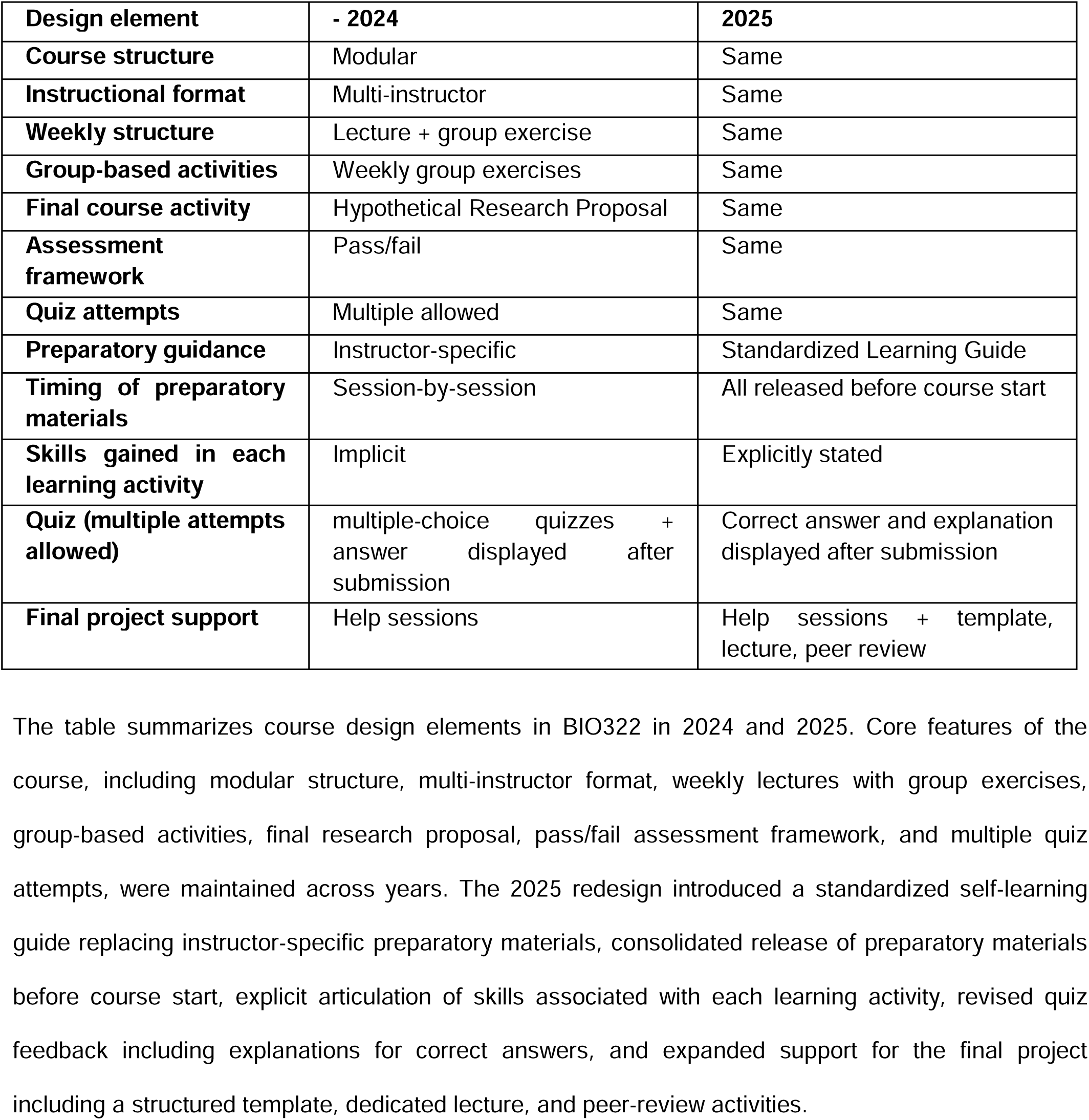
Comparison of BIO322 course design elements before (2024) and after (2025) standardization of learning support.

| Design element | - 2024 | 2025 |
| --- | --- | --- |
| Course structure | Modular | Same |
| Instructional format | Multi-instructor | Same |
| Weekly structure | Lecture + group exercise | Same |
| Group-based activities | Weekly group exercises | Same |
| Final course activity | Hypothetical Research Proposal | Same |
| Assessment framework | Pass/fail | Same |
| Quiz attempts | Multiple allowed | Same |
| Preparatory guidance | Instructor-specific | Standardized Learning Guide |
| Timing of preparatory materials | Session-by-session | All released before course start |
| Skills gained in each learning activity | Implicit | Explicitly stated |
| Quiz (multiple attempts allowed) | multiple-choice quizzes + answer displayed after submission | Correct answer and explanation displayed after submission |
| Final project support | Help sessions | Help sessions + template, lecture, peer review |

The table summarizes course design elements in BIO322 in 2024 and 2025. Core features of the course, including modular structure, multi-instructor format, weekly lectures with group exercises, group-based activities, final research proposal, pass/fail assessment framework, and multiple quiz attempts, were maintained across years. The 2025 redesign introduced a standardized self-learning guide replacing instructor-specific preparatory materials, consolidated release of preparatory materials before course start, explicit articulation of skills associated with each learning activity, revised quiz feedback including explanations for correct answers, and expanded support for the final project including a structured template, dedicated lecture, and peer-review activities.

Quiz-related support was also revised in 2025. Mandatory module-level online quizzes were updated to include brief explanations for correct responses after submission. Quiz questions were reviewed and revised to improve alignment with lecture content, module-level learning objectives, and the standardized preparatory guidance. Two types of optional quizzes were introduced: an early-stage knowledge-check quiz before the first module and a quiz linked to the final group-based research design assignment. Additional final-project support included a dedicated lecture, a structured template, worked examples, and peer-review activities. After submission of group exercises, model answers were provided newly in 2025.

### Data Sources

This study used data from completed BIO322 course iterations from 2021 to 2025. The analysis was conducted after the 2025 course had ended and used only data generated through routine course administration and institutional course evaluation.

Institutional course evaluation data were available for 2021, 2022, 2023, 2024, and 2025. These data included aggregated course evaluation scores and anonymous free-text comments collected through the university-administered evaluation system. The course evaluation responses were fully anonymous to the instructor. Institutional course evaluation response rates were 52.50% in 2021, 50.68% in 2022, 89.33% in 2023, 43.86% in 2024, and 62.67% in 2025.

Learning management system access data were available for 2023, 2024, and 2025. These data consisted of aggregated resource-level access summaries, including total numbers of accesses to course resources and numbers of students accessing each resource. These data were not available for 2021 and 2022..

The analysis therefore combined three data sources: (1) aggregated course evaluation scores (2021–2025), (2) aggregated learning management system resource-access data (2023–2025), and (3) anonymous free-text evaluation comments (2023–2025).

Free-text responses were used only to contextualize course-level patterns and identify recurring themes. They were not formally coded, and no linkage was made between comments, resource access, or assessment data. No individual-level or personally identifiable data were included or analyzed.

### Data processing and definitions

Resource-access measures were calculated from aggregated learning management system records. For each resource, the student access rate was defined as the number of students who accessed the resource divided by the total number of registered students in the course. For 2025, the denominator was 87 registered students. Page views per accessing student were calculated as the total number of page views for a resource divided by the number of students who accessed that resource.

Course resources were classified into functional categories before analysis. Resources were categorized as course navigation, course announcements, preparatory guidance, lecture slides, exercises and submissions, quizzes, final project support, further reading, model answers, or other, based on resource titles and the original learning management system category labels. Resources categorized as “other” were excluded from the category-level resource-access summary. The Course Home page was excluded from the page-views-per-accessing-student panel because it functioned as the main learning management system entry point and generated disproportionately high access counts.

### Analysis of course evaluation data

Institutional mean scores for BIO322 and faculty-wide mean scores were summarized for each evaluation item and year. For each item, the 2021–2024 range of BIO322 scores was used as a pre-2025 reference range, and 2025 scores were compared descriptively with this range, the 2021–2024 median, and the 2025 faculty-wide mean. No inferential statistical tests were applied to course evaluation scores.

### Analysis of resource access patterns

For the analysis of preparatory materials, session-linked preparatory resources from 2023 and 2024 were ordered according to their position in the course sequence. The 2025 self-learning guide was treated as a single course-wide preparatory resource and plotted at the beginning of the sequence because it was released before the start of the course and covered all teaching sessions.

For lecture slides and further-reading items, resources were ordered according to their corresponding course session. Quiz resources were classified as mandatory or optional. Module 0 and Module 4 quizzes were categorized as optional, whereas Module 1, Module 2, Module 3, and the final quiz were categorized as mandatory.

Descriptive summaries included the minimum, median, and maximum student access rate; the number of resources with student access rate above 0.5; and the minimum, median, and maximum page views per accessing student.

### Statistical software

All analyses and visualizations were conducted in R (version 4.6.0) [29]. The following packages were used: tidyverse, readr, lubridate, stringr, patchwork, and scales [30–32].

### Ethics Statement

This study was initially registered with the Norwegian Agency for Shared Services in Education and Research (Sikt; reference number 353306) for the planned processing of individual-level student data based on consent, in accordance with GDPR Article 6(1)(a). The original plan covered individual-level learning management system data and assessment-related records, with legitimate interest under GDPR Article 6(1)(f) used only to contact students for consent.

Students were invited to provide consent and a reminder was sent. Because no consent responses were received, the planned individual-level analysis was not conducted. Following consultation with institutional data protection advisors, the study design was revised to use only anonymized institutional course evaluation summaries and aggregated, non-identifiable learning management system resource-access summaries. Sikt was informed that individual-level personal data would no longer be processed for this study.

The revised study design was assessed through institutional data-protection consultation and recorded in the Sikt registration. The dataset analyzed by the authors contained only anonymized or aggregated course-level records and no identifiable human-subject data. Under these conditions, the study did not involve processing of personal data and did not require further ethics approval under the applicable institutional and data-protection procedures.

The present analysis uses only data that had been anonymized or aggregated before analysis by the authors: (1) anonymized institutional course evaluation results collected and administered by the Norwegian University of Life Sciences, and (2) aggregated resource-level summaries of learning management system access. These data contain no names, student identifiers, demographic variables, individual assessment records, or individual activity logs, and cannot be linked to individual students.

Course evaluation data were collected as part of routine institutional course evaluation procedures and were fully anonymous to the instructor. Participation in course evaluation was voluntary. No consented individual-level dataset was created, and no individual-level data were analyzed.

### Use of A Large Language Model

AI-based language models were used to support English grammar checking, stylistic refinement and R coding. All conceptual content is the authors’ own.

## Results

### Standardizing course navigation was associated with improved course evaluation scores

Institutional anonymous course evaluation scores improved in the 2025 course iteration compared with previous years **(Figure 1**). Five of the six evaluation items reached their highest observed BIO322 scores in 2025, including “understanding of learning objectives”, “course structure and organization”, “contribution of other learning activities”, “perceived learning outcome”, and “overall satisfaction”. In contrast, the contribution of lectures to learning remained within the range observed in previous BIO322 iterations, suggesting that the main change in student experience was associated with course structure navigation and explicit, accessible learning support rather than lecture delivery and course format alone.

**Figure 1.**
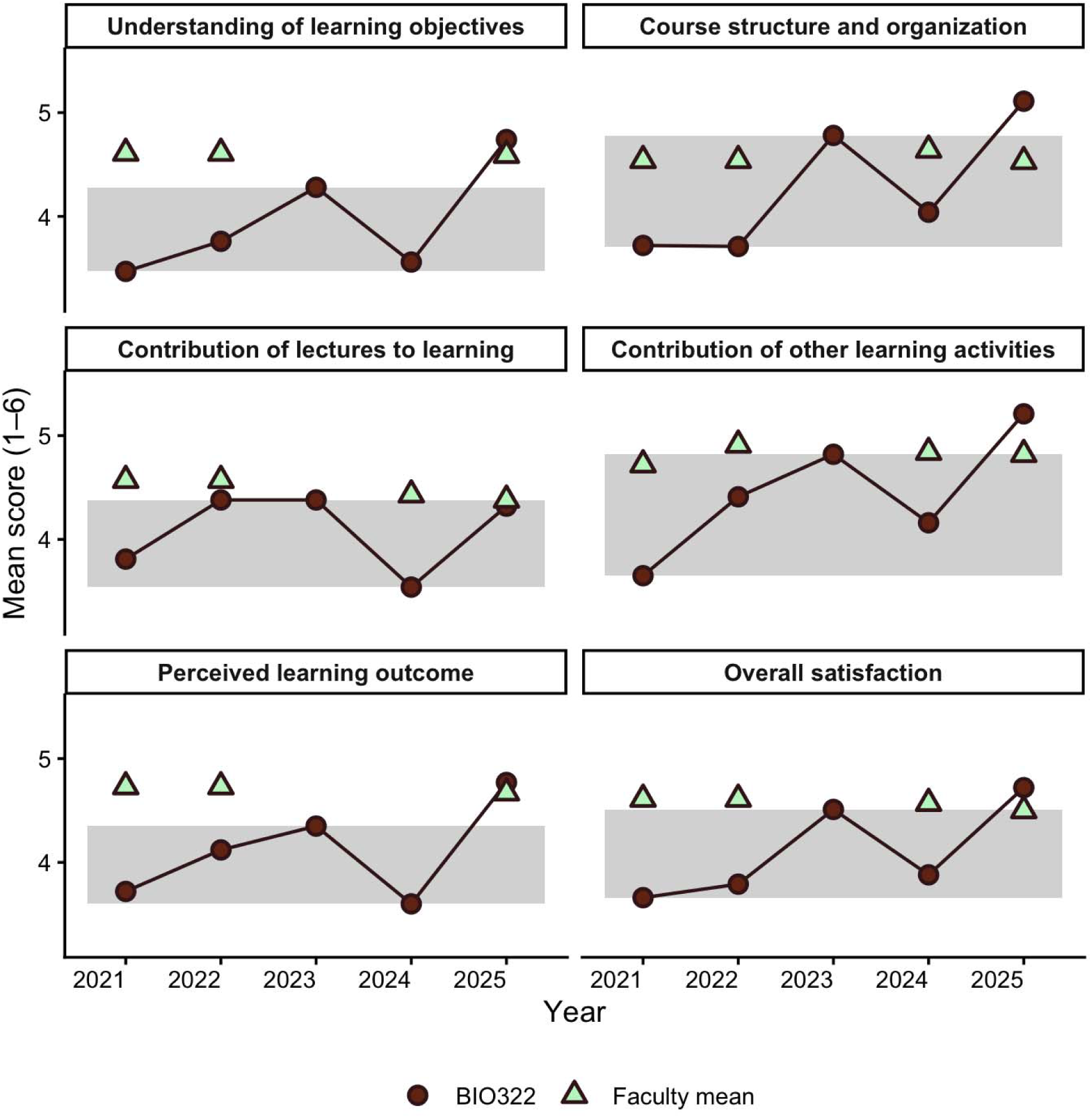
Mean course evaluation scores for BIO322 from 2021 to 2025 across six evaluation items: “understanding of learning objectives”, “course structure and organization”, “contribution of lectures to learning”, “contribution of other learning activities”, “perceived learning outcome”, and “overall satisfaction”. Circles connected by lines indicate BIO322 means by year. Triangles indicate faculty-wide means for the corresponding evaluation items in each year. Shaded regions indicate the range of BIO322 scores observed from 2021 to 2024 for each evaluation item.

### The 2025 course-wide guide replaced session-linked preparatory materials, which had been progressively less accessed in earlier years

Access to session-linked preparatory materials in 2023 and 2024 showed a decline across the course sequence (**Figure 2**). Early preparatory items were accessed by a larger proportion of registered students, whereas later items, particularly those in module 3, were accessed by fewer students. Views per accessing student also tended to be lower for later preparatory items than for early items. Across the 2023–2024 session-linked preparatory items, the median student access rate was 0.507 and the median number of page views per accessing student was 2.946. This indicates that, when preparatory materials were distributed as separate session-linked resources, both the breadth of access and repeated access among accessing students declined as the course progressed.

**Figure 2.**
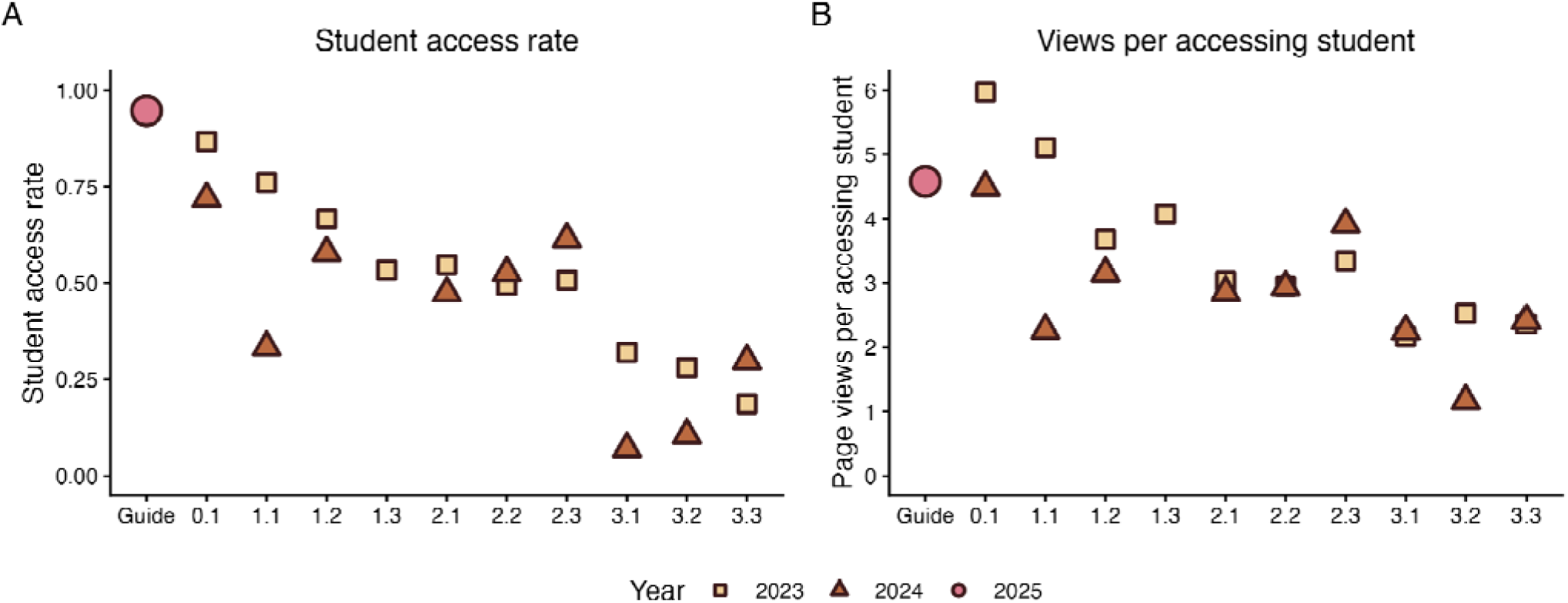
Access to preparatory materials before and after guide consolidation. Student access to preparatory materials in 2023, 2024, and 2025. (A) Proportion of registered students who accessed each preparatory content item, calculated as the number of students accessing the item divided by the total number of registered students. (B) Page views per accessing student, calculated as total page views divided by the number of students who accessed the item. The consolidated self-learning guide introduced in 2025 is plotted at the beginning of the course sequence because it was released before course start and covered all sessions. Session-linked preparatory items from 2023 and 2024 are ordered according to their position in the course sequence from 0.1 to 3.3. In 2023, the preparatory materials for sessions 1.2 and 1.3 were combined into a single item and are therefore represented at 1.2. Symbols denote course year.

The consolidated self-learning guide introduced in 2025 showed a different access pattern because it was released as a single course-wide resource before the start of the course. Nearly all registered students accessed the guide at least once. The guide had a student access rate of 0.947 and 4.577 page views per accessing student. However, views per accessing student for the guide were not clearly higher than those observed for the most frequently viewed early preparatory items in 2023 and 2024. Thus, the main observable difference was not an increase in repeated Canvas views per accessing student, but the replacement of multiple session-linked access points with one centralized resource that reached almost the entire cohort.

Because the guide was available as a single downloadable file, the access data cannot show whether students read the guide progressively across the course, downloaded it for later offline use, or accessed it only briefly. The data therefore support a limited interpretation: consolidation reduced the need for students to locate and open multiple separate preparatory files and prevented preparatory guidance from being split across progressively less-accessed session-linked resources. Whether this translated into sustained reading or preparation throughout the course cannot be determined from Canvas access logs alone.

These results suggest that consolidating preparatory guidance into a single course-wide resource made the material more visible and broadly accessed. However, because the data are aggregated at the resource level, they do not show how individual students used the guide or whether access reflected preparation before specific teaching sessions.

### Resources linked to pass/fail requirements were accessed more broadly than optional supplementary resources

To characterize how students interacted with different components of the course structure, we summarized page views and the number of students accessing resources across functional resource categories (**Figure 3**). Resources linked to course progression and assessment were accessed by larger numbers of students than optional supplementary resources. Course navigation resources and quiz resources showed high access, consistent with their direct role in locating course information and completing individual pass/fail assessment tasks. Final project support materials, exercises, and submission pages were also accessed by relatively large numbers of students, although these activities were completed in groups. Therefore, access counts for these resources may underestimate group-level use, because some students may have relied on group members to lead submission or material access.

**Figure 3.**
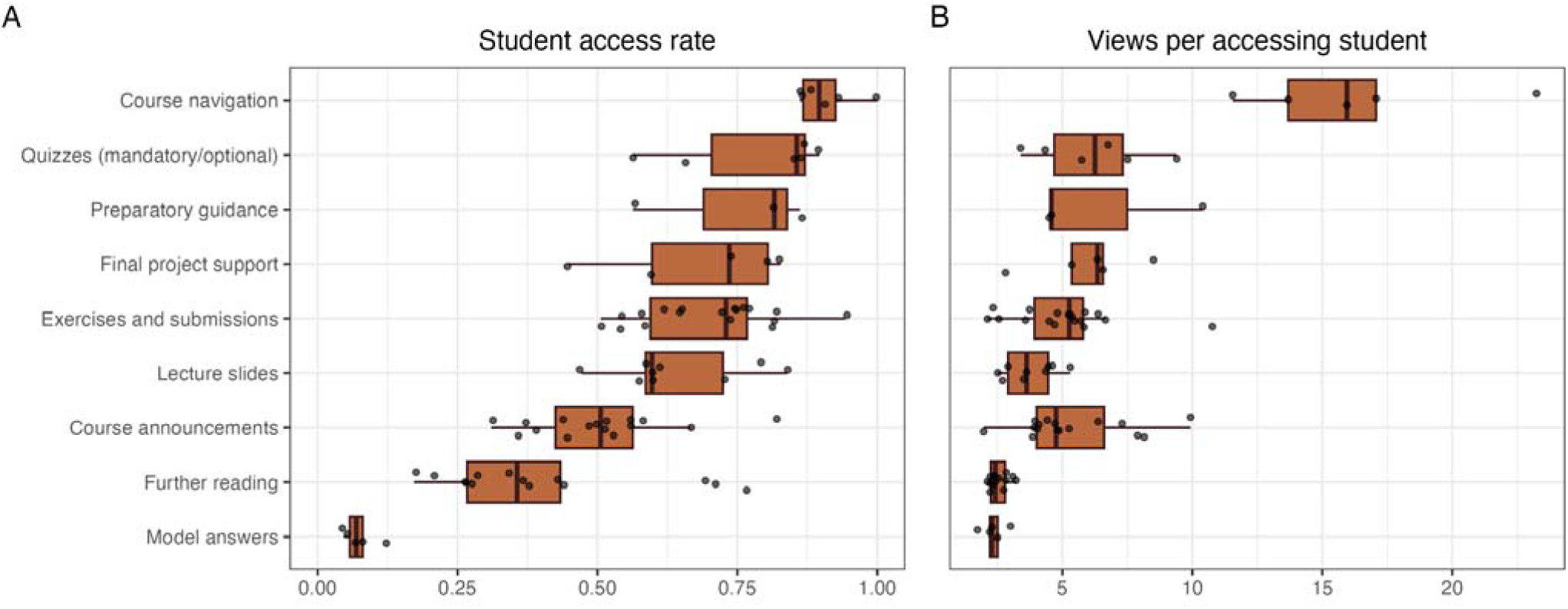
Resource access by category in the 2025 course. (A) Student access rate for each resource, calculated as the number of students accessing the resource divided by the number of registered students. (B) Page views per accessing student, calculated as total page views divided by the number of students who accessed each resource. Each point represents one course resource. Boxplots summarize the distribution within each resource category. The Course Home page was omitted from panel B because it functioned as the main Canvas entry point and generated 589.1 page views per accessing student.

Preparatory guidance and lecture slides were not themselves pass/fail requirements, but they were also accessed by relatively large numbers of students. In contrast, further-reading items and exercise model answers were accessed by fewer students. The particularly low access to model answers may reflect their position after submission: once the required task had been completed, students may have had less immediate incentive to return to model answers, especially as subsequent weekly tasks became available.

Access to lecture slides remained relatively broad across the 2025 course sequence (**Figure 4**). Although student access rates fluctuated across sessions, most lecture-slide files were accessed by more than half of registered students. Page views per accessing student also remained within a limited range across sessions. This suggests that lecture slides functioned as central course resources that students continued to access throughout the course.

**Figure 4.**
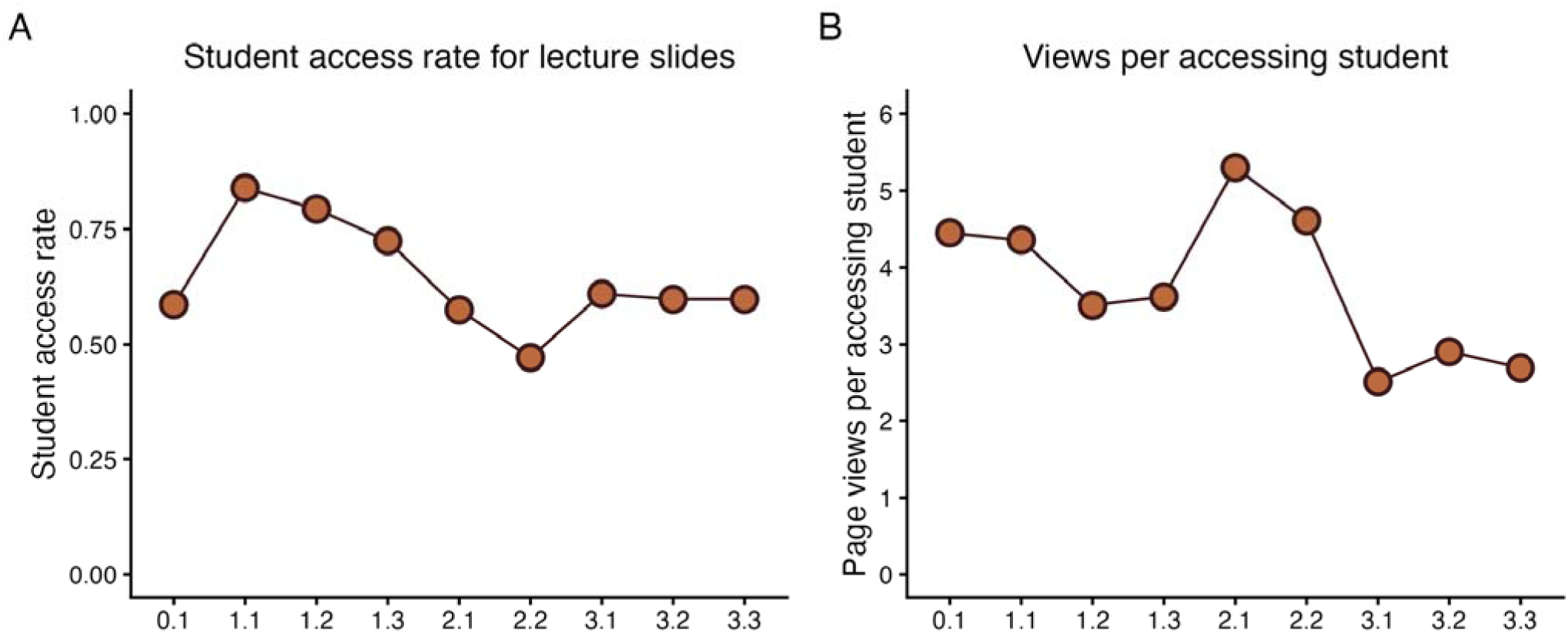
Access to lecture slides across the 2025 course sequence. (A) Student access rate for each lecture-slide file, calculated as the number of students accessing the file divided by the number of registered students. (B) Page views per accessing studnt, calculated as total page views divided by the number of students who accessed the corresponding lecture-slide file.. Lecture-slide files are ordered according to their position in the course sequence from 0.1 to 3.3.

Access to further-reading items showed a different pattern (**Figure 5**). Student access rates declined across the course sequence; Items linked to the introductory session and early Module 1 sessions were accessed by a larger proportion of students, whereas items linked to later modules were accessed by fewer students. One exception is the item labelled 4.0, which corresponded to final-project support reading. In contrast, page views per accessing student were broadly similar across further-reading items. This indicates that the main change was whether students accessed the optional readings at all, rather than how often accessing students returned to them.

**Figure 5.**
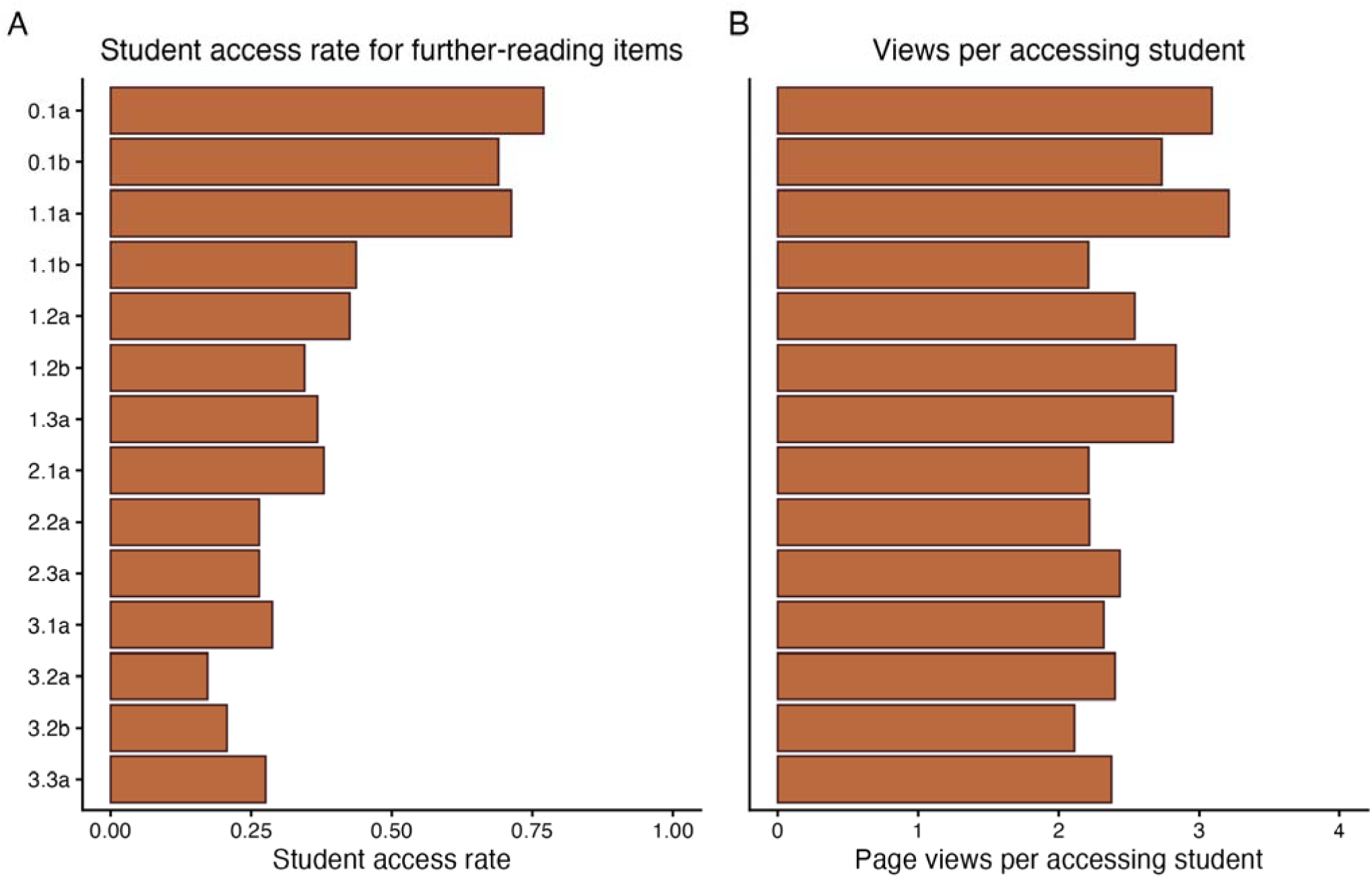
Access to further-reading items in the 2025 course. (A) Student access rate for each further-reading item, calculated as the number of students accessing the item divided by the number of registered students. (B) Page views per accessing student, calculated as total page views divided by the number of students who accessed the item. Labels indicate the corresponding course session. The item labelled 4.0 corresponds to the final-project support reading.

Together, the lecture-slide and further-reading patterns suggest a distinction between core and optional resources. Across lecture-slide files, the median student access rate was 0.598, and 8 of 9 lecture-slide files had an access rate above 0.5, while for further-reading items, the median student access rate was 0.368, and only 4 of 14 items had an access rate above 0.5. Lecture slides may have been perceived as necessary for following lectures, exercises, or assessments, whereas further-reading items may have been treated as supplementary resources. The decline in further-reading access across the course sequence may also reflect increasing workload, reduced available study time, or changing prioritization as the semester progressed. These interpretations remain tentative because the aggregated data show resource access only and do not capture student motivation, offline use, or reasons for non-access.

Quiz resources showed a third pattern (**Figure 6**). Mandatory quizzes for Modules 1–3 and the Final quiz had higher student access rates than the optional Module 0 and Module 4 quizzes. Mandatory quizzes had a median student access rate of 0.868, compared with 0.609 for optional quizzes. The median number of page views per accessing student was 7.128 for mandatory quizzes and 3.861 for optional quizzes. This pattern is consistent with the distinction between required assessment-related resources and optional support resources. Because the data are aggregated at the resource level, they describe access to quiz resources rather than quiz completion or individual assessment performance.

**Figure 6.**
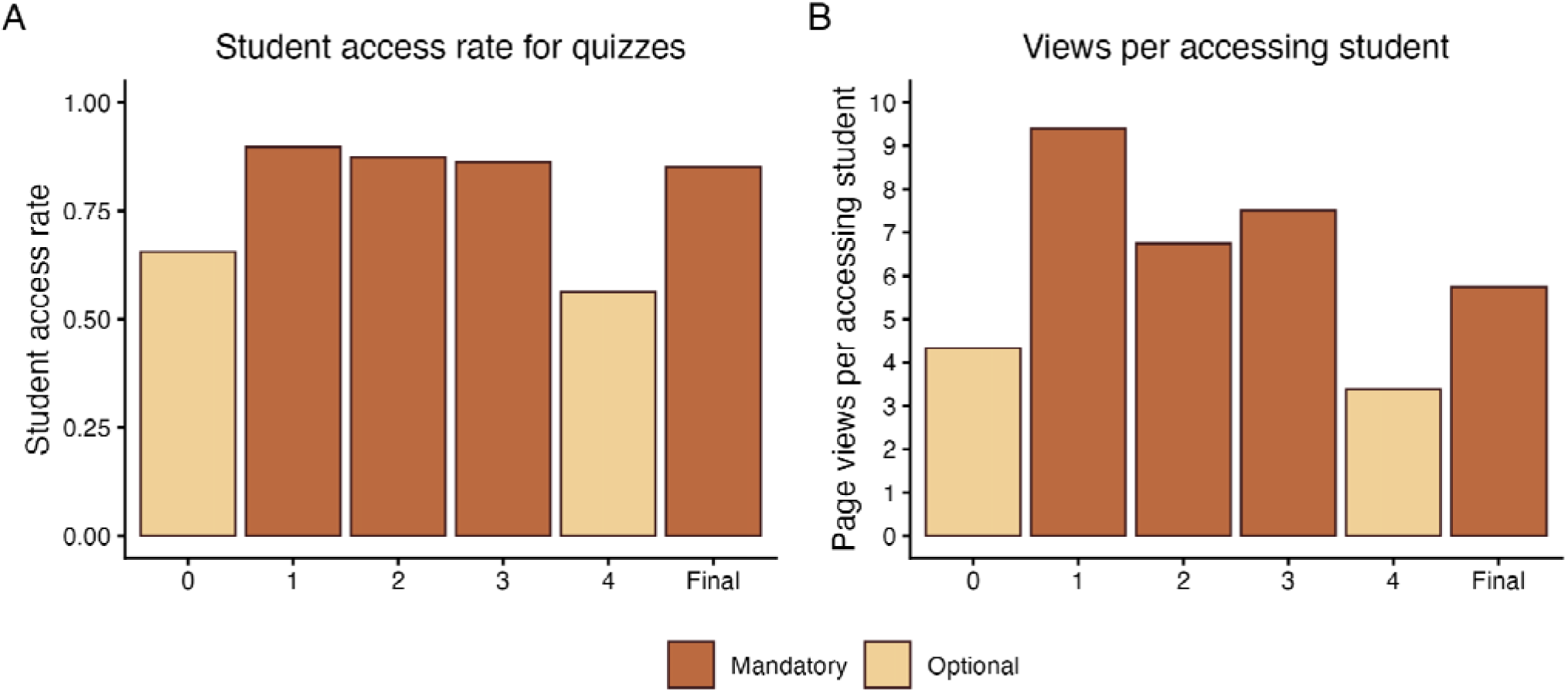
Access to quiz resources in the 2025 course. (A) Student access rate for each quiz resource, calculated as the number of students accessing the quiz resource divided by the number of registered students. (B) Page views per accessing student, calculated as total page views divided by the number of students who accessed the corresponding quiz resource. Numbers on the x-axis indicate module numbers. Module 0 and Module 4 quizzes were optional, whereas Module 1, Module 2, Module 3, and the Final quiz were mandatory.

Anonymous free-text comments from 2023–2025 contextualized the quantitative course evaluation trends. In 2023, students frequently identified weekly exercises, group assignments, lectures, feedback, and the final project as supporting learning, while also requesting clearer links between lectures and assignments and more explicit guidance for Module 4. In 2024, comments more frequently pointed to navigational problems, including unclear expectations for the semester assignment, delayed or dispersed preparatory materials, diffuse exercises, and difficulty understanding what students were expected to learn. In 2025, positive comments again emphasized weekly exercises, group work, quizzes, course structure, and the self-learning guide. However, some comments still identified remaining challenges, including information density, final-project guidance, and perceived mismatches among lecture slides, the study guide, and group assignments. These comments were consistent with the quantitative pattern of improved perceived organization in 2025, while also indicating that navigability challenges remained for some students.

## Discussion

The present study examined how a targeted redesign of course navigation was associated with changes in students’ experience and resource access in a multi-instructor graduate genomics course. Across the three research questions, the results consistently indicate that making the existing course structure more explicit was associated with measurable changes at the course level. Institutional evaluation scores improved in domains related to “understanding of learning objectives”, “course organization”, and “perceived learning outcome”, while “contribution of lectures to learning” remained stable, suggesting that the observed improvement was not driven by changes in content delivery but by increased visibility and accessibility of the course structure. Consolidating preparatory materials into a single self-learning guide released before the course began was associated with broader access across the student cohort, replacing fragmented access to session-specific materials. Resource access patterns further indicate that students selectively prioritized materials linked to course progression and assessment, while access to optional resources remained limited and declined over time. Together, these findings suggest that the intervention primarily affected how students navigated and interpreted the course, rather than altering the overall volume of engagement. These observations are consistent with prior course-design studies that evaluate implementation, student experience, and resource use as course-level outcomes, rather than treating assessment performance as the sole indicator of educational value [33,34].

### Course structure and student-perceived navigability

These results support the interpretation that, in multi-instructor graduate courses, the central challenge may lie not in the absence of instructional structure but in its accessibility from the student perspective. Prior research has shown that team-taught courses can be experienced as fragmented due to variation in instructional styles and expectations, even when the underlying structure is coherent [9,10]. In the present case, learning objectives, preparatory expectations, and course organization were consistently defined at the instructor level, yet student feedback indicated difficulty in identifying how these elements connected across sessions. This pattern is consistent with a mismatch between instructional intention and student perception, as discussed in studies of instructional accessibility [24]. Unless the relationships among objectives, activities, and assessments are made explicitly visible, alignment may exist at the design level while remaining opaque to students. This interpretation is consistent with the perspective of constructive alignment [25]. The observed improvement in evaluation items related to structure and learning objectives is consistent with the interpretation that the intervention primarily enhanced the accessibility of existing alignment rather than introducing new alignment. This also aligns with course-redesign studies in which the main pedagogical contribution lies in making expectations, structure, and learning pathways more transparent to students [35].

The findings can also be interpreted through the lens of cognitive load theory. In earlier course iterations, preparatory information was distributed across heterogeneous, instructor-specific materials, requiring students to infer priorities and relationships among topics. Such conditions may increase extraneous cognitive load by shifting effort toward interpreting course organization rather than engaging with disciplinary content [36,37]. The introduction of a standardized self-learning guide may have reduced this need for inference by explicitly signaling key concepts, expectations, and connections across sessions. While cognitive load was not directly measured, the improvement in perceived clarity and course organization is consistent with a possible reduction in extraneous processing demands. Similar effects have been reported in science education contexts, where structured preparatory materials and explicit signaling of priorities support more efficient engagement with complex content [19].

The role of preparatory scaffolding in this study appears to be primarily navigational rather than content-expanding. The self-learning guide did not introduce new material but reorganized existing content into a consistent format that supported orientation, prioritization, and connection across modules. The intervention was designed to make existing course elements more usable instead of introducing new learning tools and activities. This interpretation aligns with prior work showing that scaffolding can function by providing entry points into complex tasks and reducing ambiguity in how to engage with available resources [27,28]. However, the present results also indicate that the availability of scaffolding does not guarantee uniform uptake. Access to optional resources remained uneven, and optional quiz resources were accessed less broadly than mandatory quiz resources, suggesting that students differ in how they recognize and utilize available support. This observation is consistent with previous findings that student engagement with learning supports is shaped by differences in prior knowledge, study strategies, and perceived relevance [23].

The 2025 comments were consistent with the quantitative improvement in perceived course organization and learning support. Students frequently identified weekly exercises, group work, quizzes, assignments, and the self-learning guide as useful for learning, and some comments explicitly described the guide as providing weekly overview and structure. However, contrasting comments also remained. Some students still described the course as information-dense, requested clearer guidance for the final project, or reported mismatches among lecture slides, the study guide, and group assignments. Thus, the intervention appears to have improved perceived navigability for many students, but it did not eliminate heterogeneous experiences of coherence. This distinction is important because navigability is not only a property of the materials provided by instructors, but also of how students encounter, prioritize, and operationalize those materials during the course [38]. Meanwhile, the persistence of contrasting comments also qualifies the interpretation of learning management system access data. Broad access to the centralized guide indicates that the resource was visible and reachable, but access alone cannot show whether students used it progressively, integrated it with lectures and exercises, or found it sufficient for task initiation. Similarly, lower uptake of optional readings and optional quizzes should not be interpreted simply as lack of motivation. It may reflect workload, uncertainty about relevance, competing courses, or difficulty identifying which optional supports were worth prioritizing [39]. Future iterations should therefore focus not only on making support available, but also on positioning key supports at the moments when students need to make concrete decisions, such as selecting a final-project topic or beginning an open-ended analysis task.

### Limitation and future directions

This study should be interpreted as a course-level evaluation of instructional design rather than a direct assessment of individual learning outcomes. No individual-level linkage was available between resource access, quiz participation, assessment outcomes, or course evaluation responses. Multiple components of the course were modified in 2025, and the analysis does not allow attribution of observed changes to any single element of the redesign. In addition, learning management system access data indicate visibility and reach of resources but do not provide evidence of actual use or learning processes. Course evaluation response rates varied across years, and group-based activities may have led to underestimation of individual access to certain resources.

Similar studies have used student perceptions and course-level implementation evidence to evaluate how students experience a redesigned learning environment [40–42]. Within this scope, the findings provide evidence of changes in course navigability, clarity of expectations, and aggregate resource access with available learning supports. Improving the visibility and accessibility of course structure may enhance students’ perceived coherence of the course and influence patterns of resource access with learning resources, even in the absence of changes to core content or assessment. At the same time, the persistence of heterogeneous resource access indicates that explicit learning supports alone may not be sufficient to reach all students. These findings highlight the importance of designing not only the structure of learning materials, but also the conditions under which different students recognize, access, and act upon them in practice.

Further analysis of heterogeneity in student resource access, including individual-level variation in the use of learning supports, was originally planned but could not be conducted because no consent responses were obtained for individual-level data processing. As a result, the present study is limited to anonymized and aggregated course-level data. Future work will be required to examine how differences in student background, study strategies, and perceived relevance shape engagement with available learning supports.

In future course iterations, additional efforts may be needed to position learning supports more explicitly at points where students must make specific decisions, such as prioritizing readings, initiating open-ended analytical tasks, and selecting final project topics. Providing support at these transition points may help reduce differences in how students recognize and use available resources.

## Conclusion

This course-level evaluation shows that standardizing learning support can make an existing course structure more visible in a multi-instructor graduate genomics course. The 2025 redesign was associated with higher perceived course organization, clearer understanding of learning objectives, and broader access to centralized preparatory guidance, without changes to the core content, modular organization, or assessment framework. At the same time, uneven access to optional resources indicates that making support available is not sufficient to ensure uniform uptake. Future course development should therefore focus on placing targeted guidance at decision points where students need to translate course structure into concrete study actions, particularly for open-ended tasks such as research proposal development.

## Supporting information

SFile2

Sfile1

## Author Contribution

**Marie Saitou:** Conceptualization; methodology; investigation; data curation; project administration; writing, review and editing.

**Célian Diblasi:** Validation; resources; writing, review and editing.

## Data Availability Statement

The analysis scripts used to summarize and visualize the aggregated course-level data are available at https://github.com/mariesaitou/paper_2025-/tree/main/BIO322. The underlying course evaluation and learning management system access data are not publicly available because they derive from institutional course administration and student-related educational records. Requests for access to the underlying data will be considered in consultation with the Norwegian University of Life Sciences and Sikt (Norwegian Agency for Shared Services in Education and Research), subject to applicable institutional, legal, and data protection requirements.

## Acknowledgments

We thank the instructors who contributed to teaching BIO322 in 2025, including Thomas Nelson Harvey, Victor Boyartchuk, Prabin Sharma-Humagain, Gareth Benjamin Gillard, and Christiaan Henkel. We also thank Simen Rød Sandve, Matthew Peter Kent, and Helle Tessand Baalsrud for their contributions to the ongoing development of the course between 2022 and 2024.

We further acknowledge the support from the PPUN (Universitetspedagogikk for vitenskapelig ansatte) program, in particular Knut Omholt, Linda Helèn Godager, and Sheona Innes, for providing opportunities and perspectives for course reflection and development.

We thank Märtha Øien Felton and Julie Sivesind for assistance with establishing the legal basis for data use, and Janna Bitnes Hagen, Ingrid Roxrud, and Gisken Trøan for administrative support.

## Supporting Information

**Supplementary File 1: Self-Learning Guide (BIO322-2025)**

**Supplementary File 2: Syllabus (BIO322-2025)**

