## Supplementary material for "Making Course Structure Visible in a Multi-Instructor Graduate Genomics Course: A Course-Level Evaluation of Standardized Learning Supports": SFile2

BIO322 Advanced Topics in Genomics
Course Introduction & Student Guide

**Norwegian University of Life Sciences (NMBU)**

**Course Responsible**:
Drs. Marie Saitou, Tom Harvey

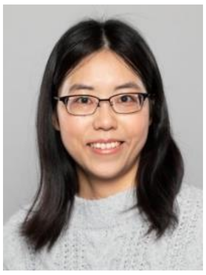 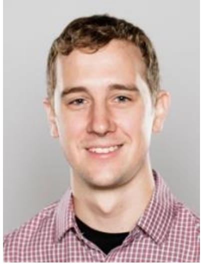

https://www.nmbu.no/om/ansatte/marie-saitou
https://www.nmbu.no/om/ansatte/thomas-nelson-harvey

#### Course Format and Schedule

**Lectures**: Mondays 10:00–12:00, Room T330
**Exercises/Group Work**: Wednesdays 08:00–10:00, Room T330

**Semester**: Autumn 2025

**Note to students: By choosing to enroll in this course, you indicate that you have reviewed the syllabus and are aware of the course requirements.**
**We encourage you to read the syllabus carefully before confirming your participation.**

**What this course is about**
BIO322 introduces how genome-scale variation drives biological differences across species, individuals, and cells.
You'll work in groups to analyze real data, explore genomic evolutionary and regulatory mechanisms and molecular and computational tools to investigate these phenomenon, and design a hypothetical research project.
Weekly tasks, quizzes, and a final proposal help you learn how to ask and answer meaningful biological questions using genomics.

### Table of Contents

*1 Overview of the Course* ... *2*

**1.1 Purpose, philosophy, and what you will learn** ... **2**

This course will help you if you: ... 2

1.1.1 Expectations for the students ... 2

**1.2 Four-module structure with progression and goals** ... **3**

**1.3 Recommended Background** ... **4**

**1.4 Inclusion and Accommodations** ... **4**

**1.5 R and Rstudio: What You're Expected to Know** ... **4**

**1.6 Language of Instruction and Support** ... **5**

*2 Module Descriptions* ... *5*

*3 Course Format and Learning Resources* ... *6*

**3.1 Instructional Methods** ... **6**

3.1.1 Lectures ... 6

3.1.2 Weekly Assignments ... 7

3.1.3 Module 4: Final Project ... 7

3.1.4 Self-Learning Tips ... 7

3.1.5 Integrated Quizzes ... 8

**3.2 Learning Resoueces** ... **9**

3.2.1 Teaching Team ... 9

3.2.2 Q&A and Feedback ... 9

**3.3 External Support Services** ... **9**

3.3.1 Writing Support ... 10

3.3.2 Statistical and Coding Support ... 10

3.3.3 Other Support Services ... 10

*4 Group Work Guidelines* ... *11*

**4.1 Group assignment policy and structure** ... **11**

**4.2 Expectations for Participation and Communication** ... **11**

**4.3 Contribution Statements** ... **11**

**4.4 Exceptional Group Reassignment and Conflict Resolution** ... **12**

*5 How to Pass This Course* ... *13*

**5.1 Assessment Components (Pass/Fail)** ... **13**

**5.2 Late Submission Policy** ... **13**

**5.3 Final Proposal Evaluation** ... **13**

**Course Schedule Summary** ... 15

### 1 Overview of the Course

#### 1.1 Purpose, philosophy, and what you will learn

BIO322 invites students to critically explore how genomic variation shapes biological diversity, from differences between species and individuals, to the regulatory mechanisms at the cellular level. Through this course, students will learn current methods and concepts in genomics, and will engage with the broader implications for environment, health, production and bioethics through genomics.

This course is grounded in the belief that genomics is not merely a technical skill set, but a powerful way to understand life**.** All living organisms share a common genetic language, and studying genomic variation allows us to trace evolutionary paths, explore functional complexity, and confront real-world issues such as biodiversity conservation, viral evolution, selective breeding and disease susceptibility.

The course promotes:

- Critical thinking based on scientific evidence
- Hands-on practice in interpreting genomic data
- Hypothesis-driven inquiry and collaborative learning
- Reflection on the societal relevance of genomics

You will learn to:

- Interpret genomic data from molecular to evolutionary scales
- Formulate and present meaningful biological questions
- Identify and justify methodological choices in genomic research
- Collaborate effectively on group projects and communicate scientific reasoning
- Engage with uncertainty and ambiguity as part of the group learning process

##### This course will help you if you:

- Are preparing for a master's thesis or career involving genomics, molecular ecology, evolutionary biology, or biotechnology
- Want to learn how evolutionary and functional genomics are used in real-world biological questions
- Need practical experience in interpreting real-world genomic data

Ultimately, our goal is to cultivate not only technical competence but also scientific literacy and responsibility, which are skills vital for contributing to society in an era shaped by fast-growing science and technology. **We hope that, through this course, you will strengthen your ability to evaluate evidence critically, consider diverse perspectives, and make informed decisions in your academic and professional life.**

##### 1.1.1 Expectations for the students

Students are expected to take an active and reflective role in their learning. This includes participating thoughtfully in discussions, cultivating self-motivation and discipline, managing their time responsibly, and seeking help when facing challenges.

If you have trouble hearing, seeing the screen, or following the lecture for any reason, please signal it during the session. Simply raising your hand is enough, and we will stop to check. If someone else has their hand raised and we have not noticed, please help by letting us know.

If speaking up is difficult for you (due to language, hearing, or communication reasons), feel free to use the a quick written message. **Please understand that if no signal is given during the session, we may not be able to address the issue later.** For example, if the room was too dim and we are told a week later by email, there is unfortunately nothing we can do retroactively. In such cases, any resulting disadvantage will be considered your own responsibility.  Thank you for helping us maintain a supportive and responsive learning environment.

#### 1.2 Four-module structure with progression and goals

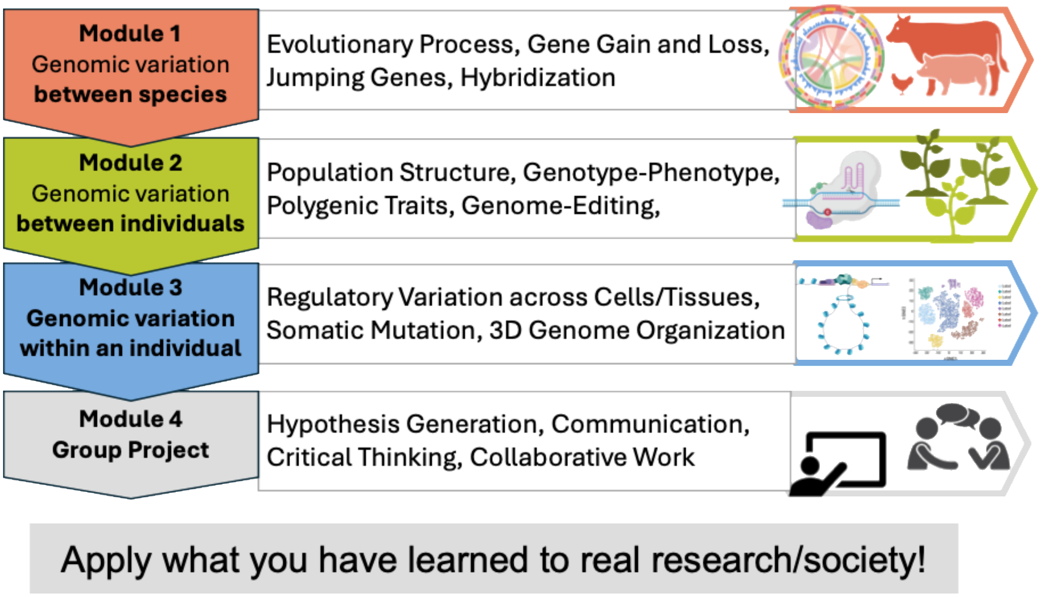


This course provides a graduate-level understanding of genome-scale variation and regulation through a four-module structure:

1. Genomic variation between species: such as between tomatoes and dogs
2. Genomic variation between individuals: such as between you and me
3. Genomic regulation within an individual: such as between liver and stomach
4. Group project: make a hypothetical research proposal

Each module targets key skills in evolutionary genomics, population genomics, and functional genomics, integrated in the group project to design a research plan. Together, these stages will help you connect genomics with real-world scientific questions.

#### 1.3 Recommended Background

This course is designed for MSc-level students in biology or related fields. You are expected to have a basic understanding of:

- Molecular biology and genetics (e.g., structure of DNA, gene expression, mutation, translation)
- Basic statistics and data interpretation (e.g., t-test, multiple testing correction)

If you feel unfamiliar with any of these topics, we recommend reviewing introductory resources.

#### 1.4 Inclusion and Accommodations

We recognize that students have diverse needs and circumstances. If you have specific challenges that may impact your participation, such as illness, disability, caregiving responsibilities, or religious events, you are encouraged to contact the course coordinator as early as possible. We will do our best to find reasonable accommodations within the scope of the course structure.

Please note:

**- Schedule conflicts with other courses are not considered grounds for accommodation.** BIO322 has a fixed schedule, and students are expected to prioritize attendance accordingly.

**- Group assignments rely on timely and consistent participation.**
If your circumstances affect your group work, please communicate clearly with both your group and the teaching team.

- **Accommodations do not guarantee deadline extensions**, but early and well-documented communication will be taken into account if issues arise.

We aim to support all students in navigating challenges constructively, while maintaining fairness and integrity in the course.

#### 1.5 R and Rstudio: What You’re Expected to Know

This course assumes basic familiarity with R and RStudio. By the end of the course, you should be able to:

- Install R packages and load them as needed
- Read and write data files from specified directories
- Produce basic plots and interpret them
- Use functions introduced in lectures or exercises (e.g., lm(), ggplot2, read.table())

You are **not expected to be an expert**, but you must be able to use R effectively for group assignments and data interpretation. If you are unfamiliar with R, please start exploaring the resources listed below before the first exercise session.

- RStudio Cloud Primers:Interactive, browser-based tutorials (no installation needed)
- Swirl: R package that teaches R inside RStudio via the console

- R for Data Science  Free online textbook (esp. Chapters 1–7 for beginners)

##### 1.6 Language of Instruction and Support

**This course is taught in English, including lecturers with diverse accents.** You are not expected to have native-level fluency, but you should be able to follow lectures, read scientific texts, and communicate your ideas in English, both in writing and in discussion.
If you feel uncertain about your English skills, we strongly encourage you to:

- Start reading the self-learning tips and core materials early
- Use the Writing Centre for feedback on clarity and structure
- Form groups with peers who are willing to support mutual understanding

**Asking for clarification during class is always welcome.** Clear communication is part of scientific training.

#### 2 Module Descriptions

**Module 1: Genomic variation between species**
In Module 1, we explore how genomes vary across species and how these differences relate to evolutionary processes. Through lectures on genome content, structure, and the effect of transposable elements (jumping genes), we learn how mutations arise, evolve and contribute to genomic and phenotypic divergence. The module also addresses mechanisms of speciation and hybridization, including the effect of genomic variation on such processes and the methods to analyze these phenomena through comparative genomics.

**Module 2: Genomic variation between individuals**
Module 2 introduces intra-species genomic diversity with a focus on the relationship between genotype and phenotype. We learn how to interpret population genetic patterns, identify signatures of evolutionary adaptation under certain environment, and understand the principles behind genome-wide association study that dissects genetic basis of phenotypic variation in a particular species (such as high or low milk yield). This module also provides an introduction to functional validation approaches, including how to test observed association signals against plausible biological mechanisms using gene-editing techniques.

**Module 3: Genomic regulation within an individual**
This module shifts the focus to within-individual regulatory variation: how identical genomes give rise to diverse cellular and functional states. We engage with transcriptomic and epigenomic data to understand both intrinsic (e.g., developmental stage) and extrinsic (e.g., environmental or mutational) sources of regulatory plasticity. Topics include somatic mosaicism, gene expression variation, chromatin structure, and 3D genome organization. The goal is to build students' ability to interpret high-dimensional functional genomics datasets in a biological context and link them to specific hypotheses about regulation, phenotype, or disease.

**Module 4: Group Project and Research Proposal**
Module 4 provides students with the opportunity to integrate what they have learned throughout the course by developing a hypothetical research proposal. Working in assigned groups, students will formulate a biologically meaningful question, outline the methods required to address it, and

justify their choices using concepts and tools from previous modules. **Each group will present their proposal in class** and receive peer and instructor feedback. **The final submission, a 2–3 page written proposal**, is expected to reflect not only scientific logic and feasibility, but also clarity of communication and responsible collaboration. This module emphasizes creativity, synthesis across disciplines, and real-world research planning and communication skills.

### 3 Course Format and Learning Resources

#### 3.1 Instructional Methods

This course combines multiple instructional methods to support conceptual understanding, practical skills, and scientific reasoning. We recognize that students come from diverse academic backgrounds and aim to provide a structured yet flexible environment for learning.

Since the field is rapidly evolving across various disciplines, there is no single textbook that covers everything comprehensively. However, as active scholars, we strive to create and update a wide range of original materials to support your learning.

All slides and data files will be shared via **Canvas**.

##### 3.1.1 Lectures

- Lectures are held weekly on Mondays.
- Instructors may vary in their teaching styles: some rely on slides, others on live explanation or diagrams.
- Slides will be shared when available, but **in-class explanations often contain key contextual material not captured in the slides.**
- Students are encouraged to ask questions during class and bring unclear points to Q&A sessions.

###### 3.1.1.1 Attendance policy and non-recorded nature of lectures

**Lectures in BIO322 are not recorded**. This is not due to pedagogical principle alone, but also because recording videos for interactive, in-person sessions is time- and resource-intensive. Instead, we focus our efforts on in-person explanation and responsive support via Canvas or after class. Thank you for your understanding and cooperation in helping us prioritize time and energy where it most benefits your learning.

**Attendance is not mandatory**, but engagement is essential. You are not penalized for missing lectures or exercises. But participation gives you access to discussion, clarification, and examples that cannot be captured in slides alone. If you miss a session, it is your responsibility to catch up. You may do so by reading the materials, contacting your group, or asking questions to the instructors. Please note that knowledge gaps due to absence may not be considered valid reason during evaluations.

**Extended Absences:** If you expect to be absent for an extended period (e.g., due to illness, family matters, or travel lasting multiple weeks), we appreciate a brief heads-up in advance.

Especially if your absence may affect your group work, early communication helps us support coordination or offer guidance.

##### 3.1.2 Weekly Assignments

Weekly exercise sessions on Wednesdays are designed to help you apply concepts from lectures to real-world data and scenarios. You will work in assigned groups to complete each task, **which must be submitted via Canvas within one week.**

While attending the in-class sessions is encouraged, it is not mandatory for all group members to be physically present. However, clear coordination and fair division of work are expected. If you cannot attend a session, please communicate with your group and ensure that your contributions support timely and complete submission.

##### 3.1.3 Module 4: Final Project

Module 4 centers around a **hypothetical research proposal**, developed and presented in groups. This project integrates knowledge from all previous modules and simulates the process of designing a real-world study.
As your final project, you will deliver a 6-minute group presentation and submit a 2–3 page written research proposal (excluding references). A template for the written report is available on Canvas.

**Attendance at the final group presentation is mandatory.**

The presentation is a core component of your course assessment and must be delivered as a group. If you anticipate being unable to attend the final presentation, you must notify the instructors in advance and discuss possible alternatives. Last-minute absences without valid reason may negatively affect your evaluation.

Students will:

- Define a biologically relevant research question
- Formulate hypotheses and propose appropriate methods to test them
- Consider the novelty and ambition of their project ideas
- Reflect on the potential academic and societal impacts of their work
- Collaborate to prepare a written proposal and group presentation

This module emphasizes critical thinking, hypothesis-driven inquiry, and scientific communication. The experience prepares students for independent research and strengthens their ability to connect genomics concepts with practical applications.

##### 3.1.4 Self-Learning Guide

To support your preparation and independent learning, you will receive short "Self-Learning Guide" for each session. These are short summaries outlining:

- Key concepts and vocabulary
- Suggested resources

Reading these tips is optional but encouraged. They are not substitutes for attending class, but they can help you get more out of the lectures and ask better questions.

##### 3.1.5 Integrated Quizzes

At the end of each module, quizzes will be available on CANVAS for review and assessment. These quizzes can be taken multiple times. **To pass, students must eventually score at least 80% on quiz 1-3 and the final quiz.**

| Skill | Activity | Lv.1. Emerging | Lv. 2. Proficient | Lv. 3 Advanced |
|---|---|---|---|---|
| 1. Understand | Lecture, Self-Learning Guide, Exercise, Quizzes | Recognizes key terms and examples when explained. | Accurately explains relevant concepts and methods introduced in the course. | Connects concepts across modules and articulates their relevance independently. |
| 2. Analyze | Exercise, Quizzes, Final Project | Identifies basic patterns or results with guidance. | Interprets data from exercises, identifies limitations, evaluates method fit. | Compares interpretations, critiques approaches, and reflects on biological logic. |
| 3. Create | Final Project | Repeats known ideas with minimal adjustments | Proposes testable questions that relate to course content. | Designs new frameworks or hypotheses drawing from multiple elements in the course. |
| 4. Communicate | Exercise, Final Project | Shares ideas when asked. Accepts group roles. | Communicates clearly in writing/presentation. Participates fairly in group work. | Leads discussion, resolves misunderstandings, and strengthens group output. |

This rubric outlines how students are expected to demonstrate understanding, analytical thinking, hypothesis generation, and collaboration throughout the course. The columns represent increasing levels of performance, from foundational to integrative.

#### 3.2 Learning Resources

##### 3.2.1 Teaching Team

- **Marie Saitou** (Course Coordinator; Lead for Modules 1 & 2)
  Responsible for course design, coordination, and teaching of Modules 1 (genomic variation between species) and 2 (variation between individuals).
  Marie is your primary contact for conceptual questions and course policies. She specializes in evolutionary genomics, focusing on genetic diversity and its evolutionary significance across species and individuals. More about her research.
- **Thomas Nelson Harvey** (Module 3 Lead & Course Coordinator)
  Leads Module 3 (regulation within individuals) and supports the Canvas usage. He applies CRISPR-based screens to identify genes linked to pathogen susceptibility and production traits. More about his research.
- **Guest lecturers and teaching contributors**
  Additional instructors will participate in specific sessions as topic specialists or co-instructors. Their contributions enhance the course's interdisciplinary depth.
- **Module 4 (Group Project)**
  Both Marie and Tom will support group project development. Questions may be directed to either instructor depending on the module background of your project.

If you are unsure whom to contact, please start with Marie or Tom. We will redirect your questions as needed.

##### 3.2.2 Q&A and Feedback

- Weekly in-person sessions are available for clarifying lecture content, assignment expectations, or conceptual questions. You can also post a question on Canvas or send an email to relevant lecturers.
- **Before asking questions, please review the syllabus** and course materials, as many common queries are addressed there.
- Students are expected to bring specific, clearly formulated questions.

#### 3.3 External Support Services

This course encourages students to develop independence and self-awareness as scientific learners. While lectures, exercises, and Q&A sessions are provided, students are also expected to make proactive use of institutional support to deepen understanding and address challenges case by case.

Seeking support is a key academic skill. Whether you are working with peers, consulting the Writing Centre, or asking for help with statistical tools, you are expected to:

- Identify what you do not understand
- Explain your issue clearly and constructively
- Revise and reflect after receiving input

We are here to support you, but we also ask that you use available resources responsibly and come prepared when asking for help.

##### 3.3.1 Writing Support

The NMBU Writing Centre provides free guidance on academic writing, including structure, clarity, and argumentation. This service is particularly valuable for improving written assignments such as group proposals.

Please note:

- You are responsible for the logic, content, and accuracy of your writing.
- If you receive writing support, you must revise the text yourself and acknowledge external input in an appendix.
- Group work must reflect your group's original reasoning and conclusions.

##### 3.3.2 Statistical and Coding Support

The Biostatistics Advising Service (BIAS group) can support you in:

- Statistical concepts and interpretation
- Troubleshooting problems in R
- Feedback on planned analytical approaches

##### 3.3.3 Other Support Services

In addition to academic support services, students may also benefit from broader institutional resources. For questions related to course planning, progression, or program changes, your study advisor is the first point of contact.
Administrative matters such as registration, transcripts, and study permits can be addressed at the Student Information Centre (SIT).

### 4 Group Work Guidelines

Group work is an essential part of this course. It is designed to help students develop collaboration skills with diverse people with different interest under real-world conditions including shared accountability. What matters most is not perfect solutions, but thoughtful engagement and fair contribution.

#### 4.1 Group assignment policy and structure

Groups are randomly assigned with three to four members, though some may have only two when some students withdraw the courses. Based on previous years, two-person groups remain viable and will not be changed. **Students may not request to select or change group members.**

If you are the sole remaining member of a group due to withdrawals or other reasons, the course team will assign you to another group (See 4.4). Reassignment decisions will be made to maintain balance and cannot be chosen by the student.

Informal collaboration across groups (e.g., sharing ideas or troubleshooting) is acceptable, as long as your final submission reflects your own group's reasoning. If relevant, clearly note such interactions in your contribution statement.

#### 4.2 Expectations for Participation and Communication

You are expected to engage consistently and communicate clearly within your group. Even if you cannot attend a weekly session in person, you are still responsible for fulfilling your part of the work and contributing to timely submission via Canvas (within one week of the session).

Group work is not about solving everything perfectly. It is entirely acceptable, and often helpful, to submit partial answer with comments such as:

- “We reached this point but didn’t fully understand,”
- “We encountered this error and couldn’t resolve it.”

This kind of comment helps instructors support your learning effectively.

#### 4.3 Contribution Statements

Every group submission must include a short contribution statement describing what each member did. This improves transparency, recognizes invisible work (e.g., coordination or editing), and helps instructors provide relevant feedback.

For example:

Odin: Wrote code for Figure 2 and ran PCA; discussed results with group
Frøya: Interpreted results and drafted paragraph 2; helped edit the file
Loki: Searched for relevant literature and proposed hypothesis; submitted final version

**If one person contributed significantly more or less,** or if external help was involved (e.g., discussion with another group), please state this briefly.

Statements will not be graded independently but may be consulted in cases of group dysfunction. If a student persistently contributes little to no work and fails to engage with the group (e.g., does not respond to messages, misses meetings), the case may be subject to follow-up.

**In such cases, the following process applies:**

- Initial reports of imbalance will prompt a group-wide check-in (not individual accusation).
- If issues persist, instructors may arrange a brief group meeting (15–20 minutes) to understand the situation.
- If a student remains non-responsive or fails to participate meaningfully, instructors may assign an alternative task (e.g., a short reflective or supplemental assignment) to ensure learning accountability.
- In extreme cases, conditional removal from the group project may be considered, with a shift to individual submission. Such decisions will be made collectively by instructors based on documented evidence (e.g., prior Contribution Statements, message history, peer comments).

We trust most students will engage constructively, but these procedures are in place to protect fairness and foster a professional and respectful learning environment.

#### 4.4 Exceptional Group Reassignment and Conflict Resolution

Group membership is assigned randomly and cannot be changed by student request. This policy reflects the reality of collaborative research, where team members are not self-selected. Differences in personality, working style, or research interest are not valid reasons for reassignment.

We ask all students to respect the trust and integrity of the group structure. Requesting to switch groups may create discomfort or unfairness for others, and will not be approved unless there is a serious, well-documented issue**.**

If you encounter a situation that makes participation genuinely difficult, such as a group member being unresponsive, or exclusion or harassment, you must:

1. **Document the issue (e.g., screenshots of unanswered messages, email attempts, communication breakdowns)**
2. Contact the instructors early via Canvas or email

Exceptional reassignment may be considered only in serious, well-documented cases (e.g. exclusion, harassment, or health-related issues). Such cases will be referred to SIT (Student Information and Support Team) for review. This may include mediation, redistribution of responsibilities, or reassignment if strictly necessary. Reassignment is not automatic, **and you may not choose which group to join.**

If a group member stops participating and cannot be reached, first attempt to resolve the issue within your group. If this fails, notify the instructors promptly with documentation. You will not be penalized for another member's silence, provided that you have communicated clearly and early.

We take group dynamics seriously and will support you in finding a fair solution, but this requires timely, clear and reasonable communication from your side.

### 5 How to Pass This Course

#### 5.1 Assessment Components (Pass/Fail)

This course is graded on a Pass/Fail basis. While the components below guide evaluation and feedback, no numerical or letter grades will be assigned.

| Component | Weight | Details |
|---|---|---|
| Weekly Group Assignments | 25% | Written responses or data interpretation tasks submitted as a group. |
| Three Module Quizzes Final Quiz | 15% | Multiple-choice format. 80% required to pass; multiple attempts allowed. |
| Final Group Proposal | 30% | 2–3 page research design + contribution report. |
| Final Presentation | 30% | 6-minute group presentation assessed on clarity and biological reasoning. |

**To pass the course**: students must (1) submit weekly assignments, (2) pass quizzes 1-3 and the final quiz, and (3) participate in the final project presentation and report. For details of each assignments, please see section 3.1.

#### 5.2 Late Submission Policy

Late submissions will be **evaluated on a case-by-case basis**, considering your **overall participation in the course**, especially your contributions to group assignments and prior engagement.
Students are encouraged to communicate any delays in advance whenever possible. Extensions may be granted in cases of illness, emergencies, or documented personal difficulties, but are not guaranteed.

Resubmission of assignments is neither required nor expected. All submissions are evaluated as-is. Use comments to guide your future work, especially the final project. Start early, review your work carefully, and seek support if needed. Exceptions (e.g., technical upload errors) are considered only with clear documentation.

#### 5.3 Final Proposal Evaluation

The final proposal will be evaluated based on the following four criteria. Each criterion will be scored on a qualitative scale (meets / partially meets / does not meet expectations). Feedback will focus on reasoning, clarity, and feasibility, not perfection.

| Criteria | Meets Expectations (✓) | Partially Meets (~) | Does Not Meet (X) |
|---|---|---|---|
| **1. Biological Relevance** | Research question is clearly stated, biologically meaningful, and logically grounded. Hypothesis is testable and coherent. | Topic is somewhat relevant but lacks clarity, specificity, or biological logic. | Topic is vague, trivial, or not related to core course themes. Hypothesis is missing or unjustified. |
| **2. Methodological Fit** | Methods are appropriate for the question, justified clearly, and reflect understanding of course concepts and tools. | Methods are partially appropriate or only weakly justified. Some misunderstanding of tools or gaps in reasoning. | Methods are inappropriate, missing, or disconnected from the research question. |
| **3. Interpretation & Limitations** | Expected outcomes are explained with insight. Potential limitations and uncertainties are acknowledged thoughtfully. | Some interpretation is present but lacks depth, balance, or awareness of limitations. | Interpretation is missing, superficial, or overly speculative. No discussion of limitations. |
| **4. Structure & Communication** | Proposal or presentation is clearly organized, concise, and persuasive. Visuals and formatting support understanding. | Some parts are unclear, redundant, or disorganized. Visuals (if used) are underdeveloped. | Proposal/presentation is hard to follow. Poor structure or confusing language. Visuals (if present) hinder clarity. |

Use this guide to understand how your project work will be assessed. Each criterion is rated as "Meets Expectations (✓)," "Partially Meets (~)," or "Does Not Meet (X)." Clarity, relevance, and critical thinking are essential.

Plagiarism, fabrication, or template reuse without acknowledgment will result in disqualification. All group members will receive the same grade unless a formal request for individual differentiation is submitted.

Especially, AI may produce fabricated references or inaccurate claims. Always verify sources and data independently. All ideas and text submitted must reflect your own understanding. Do not submit AI-generated content that you do not fully comprehend or cannot justify.

#### Course Schedule Summary

| Uke | Ukedag | Date | Time | Session ID | Topic | Responsible | Deadline |
|---|---|---|---|---|---|---|---|
| u 36 | Onsdag | 03.09.2025 | 08:00–10:00 | 0 | Introduction | Marie | |
| u 37 | Mandag | 08.09.2025 | 10:00–12:00 | 1.1 Lecture | Genome evolution | Marie | |
| u 37 | Onsdag | 10.09.2025 | 08:00–10:00 | 1.1 Exercise | Genome evolution | Marie | |
| u 38 | Mandag | 15.09.2025 | 10:00–12:00 | 1.2 Lecture | Transposable Elements | Marie | |
| u 38 | Onsdag | 17.09.2025 | 08:00–10:00 | 1.2 Exercise | Transposable Elements | Marie | 1.1 Exercise |
| u 39 | Mandag | 22.09.2025 | 10:00–12:00 | 1.3 Lecture | Speciation/Hybridization | Marie | |
| u 39 | Onsdag | 24.09.2025 | 08:00–10:00 | 1.3 Exercise | Speciation/Hybridization | Marie | 1.2 Exercise |
| u 40 | Mandag | 29.09.2025 | | Høstferie | | | |
| u 40 | Onsdag | 01.10.2025 | | Høstferie | | | 1.3 Exercise, Quiz 1 due |
| u 41 | Mandag | 06.10.2025 | 10:00–12:00 | 2.1 Lecture | Population Genomics | Marie | |
| u 41 | Onsdag | 08.10.2025 | 08:00–10:00 | 2.1 Exercise | Population Genomics | Marie | |
| u 42 | Mandag | 13.10.2025 | 10:00–12:00 | 2.2 Lecture | GWAS & eQTL | Marie | |
| u 42 | Onsdag | 15.10.2025 | 08:00–10:00 | 2.2 Exercise | GWAS & eQTL | Marie | 2.1 Exercise |
| u 43 | Mandag | 20.10.2025 | 10:00–12:00 | 2.3 Lecture | Genome Editing | Tom | |
| u 43 | Onsdag | 22.10.2025 | 08:00–10:00 | 2.3 Exercise | Genome Editing | Tom | 2.2 Exercise |
| u 44 | Mandag | 27.10.2025 | 10:00–12:00 | 4. Final project | Research Proposal Q&A | Marie + Tom | |
| u 44 | Onsdag | 29.10.2025 | 08:00–10:00 | 4. Final project | Research Proposal Q&A | Marie + Tom | 2.3 Exercise, Quiz 2 due |
| u 45 | Mandag | 03.11.2025 | 10:00–12:00 | 3.1 Lecture | Current Transcriptomics | Tom/Guest | |
| u 45 | Onsdag | 05.11.2025 | 08:00–10:00 | 3.1 Exercise | Current Transcriptomics | Tom/Guest | |
| u 46 | Mandag | 10.11.2025 | 10:00–12:00 | 3.2 Lecture | Somatic Mutation | Tom/Guest | |
| u 46 | Onsdag | 12.11.2025 | 08:00–10:00 | 3.2 Exercise | Somatic Mutation | Tom/Guest | 3.1 Exercise |
| u 47 | Mandag | 17.11.2025 | 10:00–12:00 | 3.3 Lecture | ATAC-seq, ChIP-seq, Hi-C | Tom/Guest | |
| u 47 | Onsdag | 19.11.2025 | 08:00–10:00 | 3.3 Exercise | ATAC-seq, ChIP-seq, Hi-C | Tom/Guest | 3.2 Exercise |
| u 48 | Mandag | 24.11.2025 | 10:00–12:00 | 4. Final project | Research Proposal Q&A | Marie + Tom | |
| u 48 | Onsdag | 26.11.2025 | 08:00–10:00 | 4. Final project | Research Proposal Q&A | Marie + Tom | 3.3 Exercise, Quiz 3 due |
| u 49 | Mandag | 01.12.2025 | 10:00–12:00 | 4. Final project | Final Presentations | Marie + Tom | |
| u 49 | Onsdag | 03.12.2025 | 08:00–10:00 | 4. Final project | Final Presentations | Marie + Tom | |
| | | 10.12.2025 | | | | | **Final proposal/Final quiz** |
