## Supplementary material for "Making Course Structure Visible in a Multi-Instructor Graduate Genomics Course: A Course-Level Evaluation of Standardized Learning Supports": Sfile1

### BIO322 Self-Learning Guide (2025)

Bridging Genomic Methods and Real-World Questions

**Course: BIO322 – Advanced Topics in Genomics**
Semester: Autumn 2025
Department: Faculty of Biosciences,
Norwegian University of Life Sciences (NMBU)

**Course Coordinators:**
Dr. Marie Saitou (Modules 1 & 2)
Dr. Thomas Nelson Harvey (Module 3)
Contact:

Lectures: Mondays 10:00–12:00, Room T330
Exercises / Group Work: Wednesdays 08:00–10:00, Room T330

### How to Use This Self-Learning Guide

This guide is not meant to be mastered in advance. Instead, it is designed to make lectures more accessible, engaging, and intellectually rewarding. Think of it as a map rather than a manual—it won't give you every answer, but it will help you orient yourself, anticipate what's coming, and recognize which questions matter.

Each session's guide includes:

- **What This Lecture Covers**
- **Vocabulary**
- **Find the Answers to These Questions During the Lecture**
- **Background Concepts** – ideas that support deeper understanding, but may not be intuitive at first
- **Recommended Resources** – selected further readings if you are interested, but reading these does not substitute attending lectures

You are encouraged to skim the guide before each lecture, bring it with you to class, and refer back to it afterward as a tool for clarification and review. You can also use it individually **or as part of a group,** discuss unfamiliar terms, think through the guiding questions together, and reflect on which topics interest you most. These group conversations may even **help shape your ideas for the Final Project: Research Proposal and Presentation.**

Also, recommended references for each session (**available on Canvas**) are selected to provide conceptual overviews. Again, you are not expected to master every method or term, but to grasp the broader questions, frameworks, and strategies presented, especially if you are interested in that particular topic. Focus on the structure of arguments, the way scientific evidence is integrated, and how genomics connects to broader biological or societal themes.

### Biological Scale in Genomics: Phenomena, Methods, and Relevance

BIO322 is designed to guide you through the different scales at which genomic variation can be observed and interpreted. In **Module 1**, we begin at the species level, where evolutionary history leaves signatures in genome size, gene content, and genome architecture. In **Module 2**, we move into populations and individuals, where ongoing processes like local adaptation, genomic basis of phenotypic diversity, and demographic history shape the genetic landscape within species. **Module 3** will later bring the focus inward, examining how a single genome gives rise to diverse cellular and regulatory states within one individual.

The examples below illustrate biological phenomena at different scales and the methods used to study them. This list is not exhaustive, but it is meant to provide an starting point.

##### Between Species

**What makes differences between species?**
Macroscopic differences between species like reptiles and mammals, such as skin type, thermoregulation, or body design, stem from lineage-specific changes in genome architecture. Comparative genomics reveals how features like hair, endothermy, and placentation emerged through gene duplication, enhancer evolution, or rewiring of developmental networks, and so on.

**Convergent evolution**
Across distantly related species, similar traits may evolve independently. Genomic analysis can detect convergent amino acid substitutions or regulatory changes, revealing underlining genetic basis. This informs our understanding of evolutionary constraints and adaptive landscapes.

**Viral evolution**
Viruses show rapid mutation. Comparative genomics and phylogenetics uncover how viruses adapt to new hosts, acquire toxicity, escape immunity, or gain new functions. This is essential for epidemiological forecasting and pandemic response.

#### Between Individuals

**Local adaptation**
Why do Tibetan highlanders thrive in thin air, or Arctic foxes stay warm in freezing cold? Genome scans can detect environment-associated variants that may affect fitness, the ability to survive and reproduce in a given environment. For example, certain variants help improve oxygen transport at high altitude or regulate fur thickness in cold climates. These insights are essential for understanding the genomic basis of how populations adapt locally and for guiding biodiversity conservation under climate change.

**Phenotypic diversity**
Human traits including height, skin color, or heart disease risk are influenced by many genes, each contributing a small effect. These are called polygenic traits (complex traits). Genome-wide association studies (GWAS) identify variants linked to such traits. By utilizing genomics, researchers can investigate genomic variants with likely biological effects, or variants that may cause disease.

**Demographic history**
Why do certain genetic disorders occur more often in particular populations? Comparative genomics helps reconstruct population histories. These include past migrations, bottlenecks (sharp reductions in population size), and population isolation events. Understanding this context is crucial for biodiversity conservation, understanding evolutionary history, and designing equitable biomedical research for diverse human populations.

**Pathogen–host coevolution**
Why are some people naturally resistant to certain infectious diseases, or some crops immune to plant viruses? Pathogens and hosts are in constant evolutionary competition. Genomics can detangle the molecular basis of this co-evolution, such as viral entry proteins versus host immune system receptors,  informing strategies in medicine and agriculture.

#### Within Individuals

**Aging**
Why do cells in older bodies behave differently from younger bodies, even though their DNA is unchanged? As we age, random mutations accumulate, epigenetic regulation weakens, and gene expression becomes less coordinated. Single-cell RNA-seq and methylation profiling reveal this

molecular disorganization, which underlies aging-associated diseases, and may suggest targets for intervention.

**Cancer development**
How does a normal cell become a tumor? Cancer develops through clonal evolution: somatic cells acquire mutations, some of which give extreme growth advantages, allowing those clones to dominate. Genomic instability accelerates this process. Techniques like mutational signature analysis and chromatin profiling identify cancer driver mutations and track tumor progression at the molecular scale, guiding early detection and personalized treatment.

**Organ differentiation**
How can a skin cell and a liver cell behave so differently, despite identical genomes? The answer lies in gene regulation. During development, chromatin becomes selectively open or closed in different cell types. Techniques such as ATAC-seq and ChIP-seq reveal these changes, helping us understand how organs form and how cellular identity is established, which is critical knowledge for regenerative medicine and developmental biology.

**Stress responses**
How do cells react to starvation, toxins, or infections? Stress triggers rapid shifts in gene expression, chromatin accessibility, and even genome organization. Using time-resolved RNA-seq and Hi-C, researchers can observe how cells reprogram themselves in real time. These mechanisms explain how organisms cope with challenges, and why stress sometimes leads to disease.

### 0.1 How We Study Genomes: Sequencing, Epigenetics, and 3D Genomics

#### What This Lecture Covers

In this lecture, we will explore the main techniques used to study genomes. It covers how DNA sequencing is performed, the difference between short-read and long-read platforms, and how sequences are assembled into larger units. It also introduces RNA sequencing and methods for detecting epigenetic modifications like DNA methylation. Finally, it explains how to study chromatin structure and the 3D arrangement of the genome using tools such as ATAC-seq, ChIP-seq, and Hi-C.

#### Vocabulary

1. **Sanger sequencing:** First-generation sequencing method; high accuracy, low throughput, moderate read length
2. **Illumina sequencing:** High-throughput, short-read sequencing platform with high accuracy
3. **Pacific Biosciences (PacBio, SMRT):** Long-read sequencing technology; moderate accuracy; useful for resolving repetitive regions and genome assembly
4. **Oxford Nanopore sequencing:** Real-time, ultra-long-read sequencing; relatively lower accuracy
5. **DNA microarray:** Hybridization-based method for detecting known single nucleotide polymorphisms (SNPs)
6. **RNA sequencing (RNA-seq):** Sequencing of RNA transcripts to investigate gene expression and transcript structure
7. **Bisulfite sequencing (BS-seq):** Method to detect cytosine methylation by converting unmethylated cytosines to uracils
8. **Assay for Transposase-Accessible Chromatin using sequencing (ATAC-seq):** Method to assess chromatin accessibility by inserting sequencing adapters into open chromatin regions
9. **Chromatin Immunoprecipitation sequencing (ChIP-seq**): Technique to identify DNA regions bound by specific proteins such as transcription factors or histones
10. **Hi-C:** Genome-wide method for capturing three-dimensional genome organization by sequencing spatially proximal DNA fragments

#### Find the Answers to These Questions During the Lecture

1. What are the trade-offs between short-read and long-read sequencing approaches?
2. How do single nucleotide polymorphism arrays (SNP arrays) work?
3. What are the main steps in preparing an RNA-seq or ChIP-seq library?
4. What is the difference between chromatin accessibility (e.g., ATAC-seq) and DNA–protein interaction mapping (e.g., ChIP-seq)?
5. How do you measure a functional unit of chromosome architecture?

#### Background Concepts

- **What is a “read” in sequencing?**
  A read is a fragment of DNA or RNA whose sequence has been determined.

- **Why does “assembly” matter?**
  Sequencing technologies generate many short fragments rather than an intact genome. These fragments must be assembled into contigs (continuous sequences), which are then organized into scaffolds and eventually complete chromosomes.

- **Gene regulation**

  Gene regulation refers to the processes that control when, where, and how much a gene is expressed in a body. For a gene to be expressed, a key step is the binding of **RNA polymerase II** to a region of DNA called the **promoter**, which is located just upstream of the gene. The promoter serves as the main entry point for the transcription machinery and determines the start site for RNA synthesis. In addition to promoters, there are other **regulatory elements** that influence gene expression. **Enhancers** are DNA regions, often located far from the gene they control, that boost transcription when bound by specific **transcription factors**. They help activate genes in the right cells at the right time. In contrast, **silencers** are regions that repress transcription when bound by **repressor** proteins.

- **Genome organization**

  In the nucleus, the genome is not stretched out like a straight line, it is folded in complex ways so that distant parts of the DNA can be close physical proximity. The contact frequency between different genomic regions reflects how often they are near each other in three-dimensional space. These contacts play important roles in **gene regulation**, **chromatin accessibility**, and **genome organization**. For example, an **enhancer** may activate a gene by physically looping over to its **promoter**, even if they are far apart along the linear genome. Likewise, regions that are frequently in contact may share similar epigenetic states. Therefore, measuring chromatin contact frequencies helps us understand how the 3D structure of the genome contributes to its function, how spatial folding influences which genes are turned on or off.

#### Recommended Resource

**Hu et al. (2021) – Next-generation sequencing technologies: An overview**
A concise guide to major sequencing platforms and their features (e.g., read length, error profiles), with deeper platform-specific chemistry reserved for advanced reading.

**Theissinger et al. (2023) – How genomics can help biodiversity conservation**
A timely and applied review that shows how a variety of genomic tools support species monitoring and ecosystem management. You can focus on main method categories.

### 1.1 Genome Size, Content, and How They Evolve

#### What This Lecture Covers

In this lecture, we will look at how genome size and gene content vary between species. It explains the processes that add or remove genes, such as genome duplication. It also introduces tools used to compare genomes between species. The lecture gives examples of how genes help organisms adapt to specific environments and why similar organisms may have very different genomes.

#### Vocabulary

1. Genome size: Total amount of DNA in the haploid genome
2. Genome architecture: The arrangement of elements within the genome
3. Segmental duplication: Duplication of specific regions of the genome, sometimes generating additional gene copies
4. *De novo* gene: A gene that originates from previously non-genic DNA sequences
5. Pseudogenization: The process by which a functional gene becomes inactive due to mutations
6. Homologous genes: genes passed down from the common ancestor of two species
7. Orthologs: Homologous genes found in different species that diverged through speciation
8. Paralogs: Homologous genes within the same species that arose via gene duplication
9. Whole-genome duplication: A process where the entire genome is duplicated
10. Rediploidization: The evolutionary process by which a polyploid genome (four set of genomes, for example) returns to a functionally diploid (two set of genomes) state

#### Find the Answers to These Questions During the Lecture

1. Why do genome sizes differ so widely among species?
2. Why does the distinction between orthologs and paralogs matter?
3. What typically happens to duplicate genes after a whole-genome duplication event?
4. How does gene duplication facilitate adaptation to extreme environments?
5. What challenges are faced when inferring gene evolutionary history across distantly related species?

#### Background Concepts

**The process of molecular evolution**
When a mutation (a change in the DNA sequence) occurs in an organism, it creates a new version of a gene, known as an **allele**. Most mutations do not change the organism's traits in any noticeable way; these are called **neutral** mutations because they neither help nor harm the organism's chances of survival or reproduction. Some mutations, however, are **deleterious**, meaning they reduce the organism's **fitness**, its ability to survive and produce offspring (such as a mutation causes disease). On the other hand, if a mutation improves fitness, it is considered **adaptive** (such as a mutation leads to cold tolerance). Over generations, **natural selection** tends to remove deleterious alleles and increase the frequency of adaptive ones in a population. This process shapes the genetic makeup of species over time, even though most mutations remain neutral and simply random **drift** without strong selection.

**Meiotic drive**

**Meiotic drive** describes cases in which some genetic elements are transmitted to offspring more frequently than would be expected under standard **Mendelian inheritance**. During **meiosis**, each chromosome (or **allele**) is typically expected to have a 50% chance of being included in a gamete. However, certain chromosomal regions, such as **centromeres**, can influence the process in a way that increases their likelihood of being passed on. This unequal transmission is known as meiotic drive. A specific example is **centromere drive**, where some centromeres bias chromosome segregation during egg formation. Although such variants may reduce the organism's overall **fitness**, they can still spread in a population due to this transmission advantage. Understanding meiotic drive helps explain why certain chromosome structures, including harmful rearrangements, become fixed in natural populations.

#### Recommended Resource

**Stange, Barrett & Hendry (2021) – The importance of genomic variation for biodiversity, ecosystems and people**

This paper links genomic variation to ecosystem function and societal outcomes, offering a broad, interdisciplinary view that combines genetics, conservation, and policy. It is suitable for students exploring the societal relevance of genomics. You can focus on understanding how intraspecific variation affects ecological processes.

**Koonin & Wolf (2010) – Constraints and plasticity in genome and molecular-phenome evolution**

This review explores the multiple levels at which evolutionary constraints act, from nucleotide sequences to regulatory networks and organismal phenotypes.

### 1.2 Transposable Elements (TEs) : Jumping Genes in a Genome

#### What This Lecture Covers

This lecture focuses on transposable elements (TEs), which are short pieces of DNA that can move within the genome. In this lecture, we will learn about different types of TEs, how they move, and how they affect genome size and structure. It also describes how some TEs are harmful or neutral, while others are reused by the host genome for new functions. The lecture also includes how TEs are studied using lab and computational methods.

#### Vocabulary

1. **Transposable elements (TEs)**: DNA sequences capable of changing their position in the genome; often referred to as "jumping genes".
2. **Retrotransposons (Class I)**: Copy-and-paste elements that use an RNA intermediate and reverse transcription (e.g., LINEs, LTRs).
3. **DNA transposons (Class II)**: Cut-and-paste elements that move via a DNA intermediate, typically using a transposase.
4. **Autonomous vs. non-autonomous TEs**: Autonomous TEs encode the enzymes required for their own movement; non-autonomous rely on others.
5. **Accordion model**: A model of genome size evolution where TE accumulation is counterbalanced by host sequence loss.
6. **Genome rearrangement**: Structural changes like duplications, deletions, or inversions often facilitated by repetitive TE sequences.
7. **TE divergence**: The accumulation of mutations in TE copies over time, leading to classification challenges.
8. **Epigenetic silencing**: Mechanisms like DNA methylation and piRNA pathways that repress TE activity.
9. **Gene regulation by TEs**: TEs can donate regulatory elements, disrupt genes, or affect chromatin state and transcription.
10. **TE domestication**: The evolutionary process by which TE-derived sequences are co-opted for host functions, such as novel genes or regulatory elements.

#### Find the Answers to These Questions During the Lecture

1. How can TEs contribute to both genome expansion and gene loss?
2. What makes some TEs harder to detect than others in genomic data?
3. How do the two main classes of transposable elements differ in mechanism?
4. In what ways do host genomes defend themselves against TE activity?
5. Why are some TE families more successful than others across lineages?

#### Background Concepts

**What does it mean to call TEs "selfish genes"?**
Transposable elements (TEs) are sequences that can increase their copy number within a host genome, regardless of their effect on the organism's condition. In some cases, their activity can be

neutral or even harmful. The term “selfish” is used within the gene-centered framework of evolution to describe such elements, those that persist and spread due to their own transmission dynamics in the host genome, rather than because they contribute to organismal function. This contrasts with host genes that tend to be retained through natural selection based on their benefit to the whole organism.

**What is the relationship between TEs and their host genomes?**
TEs are embedded within the host genome, yet their presence introduces a dynamic interaction over evolutionary time. While TEs can increase in number through mechanisms like transposition, host organisms have evolved molecular pathways that limit or silence TE activity. This reciprocal pressure, often described as an evolutionary arms race between TE and their host, can influence host genome structure, regulation, and gene evolution. In some cases, sequences derived from TEs are later maintained and integrated into host functions, a process known as **TE domestication.**

**How can mutation accumulation be used to estimate TE age?**
After a TE inserts into the genome, it begins to accumulate mutations over time. Mutations tend to accumulate at relatively stable rates over time. Therefore, the degree of sequence divergence from the inferred **ancestral (consensus) sequence** can be used to estimate how long ago the insertion occurred. A group of nearly identical copies suggests a recent transposition event, whereas more divergent copies reflect ancient transposition events.

**Are all TEs the same across species?**
Not at all. While the basic mechanisms (e.g., retrotransposition or DNA transposition) are shared, the actual TE families present, their structure, their regulatory environments, and their evolutionary histories are highly lineage-specific. Even closely related species can differ greatly in their TE composition, activity, and genomic impact. Comparative genomics shows that each lineage has developed a distinct TE landscape, shaped by its own evolutionary path.

#### Recommended Resource

**Bourque et al. (2018) – Ten things you should know about transposable elements**
This comprehensive review introduces the major roles of transposable elements (TEs) in genome structure, function, and evolution. It explains why they present both challenges and opportunities in genomic research..

**Almeida et al. (2022) – Taming transposable elements in vertebrates: from epigenetic silencing to domestication**
This paper provides a structured overview of how vertebrates control transposable elements (TEs) through epigenetic silencing and how some of these elements are later co-opted to serve beneficial functions.

### 1.3 Hybridization and Speciation: How Do New Species Emerge?

This lecture explains how new species form and what genetic changes are involved in that process. It introduces different types of speciation processes based on geographic context and describes barriers to reproduction that keep populations separate. It shows how hybridization can mix genes between species and sometimes create new species. The lecture also explains how structural changes in chromosomes can help or prevent gene flow between populations.

#### Vocabulary

1. **Speciation**: The evolutionary process through which new, reproductively isolated species arise from existing populations.
2. **Introgression** : The movement of genes from one species into another via hybridization and backcrossing. Detectable via methods like the ABBA-BABA test.
3. **Reproductive isolation** : Barriers preventing gene flow between populations, enabling the process of speciation.
4. **Hybrid zone** : A geographic region where genetically distinct populations come into contact and interbreed, often producing hybrids of varying fitness.
5. **Assortative mating** : A prezygotic mechanism where individuals preferentially mate with similar phenotypes, reinforcing reproductive isolation.
6. **Allopatric / Sympatric / Para-patric speciation** : Speciation that occurs with no / full / partial geographic overlap between diverging populations.
7. **Prezygotic / Postzygotic barriers** : Mechanisms of isolation that act before vs after fertilization, such as mate choice (prezygotic) or hybrid sterility (postzygotic).
8. **Incompatibility loci** : Genes that cause reduced fitness in hybrids due to negative interactions. Often evolve rapidly and can act as speciation genes.
9. **Genetic hitchhiking** : The process by which neutral or weakly selected loci increase in divergence because they are linked to loci under strong selection.
10. **Standing genetic variation** : Pre-existing genetic diversity within a population that can be utilized for rapid adaptation to environmental changes.

#### Find the Answers to These Questions During the Lecture

1. What processes initiate reproductive isolation between populations?
2. Why do some genomic regions show higher differentiation than others?
3. Why is standing variation more important than new mutations in rapid speciation?
4. How can we distinguish between differentiation due to selection vs demographic history?
5. How can chromosomal rearrangements such as inversions and fusions promote speciation?

#### Background Concepts

**Is Speciation a Binary Switch?**
"Speciation" may sound like a sudden event, either two groups are different species or they are not. But in reality, speciation is a gradual, long-term process.

Imagine a grass species that grows both on exposed mountain slopes and in shaded valley bottoms. Over evolutionary timescale, slope populations adapt to harsh, dry conditions with waxy leaves and early flowering by accumulating mutations, while valley populations evolve broad leaves and flower later in cooler, wetter habitats by accumulating other mutations. As these differences pile ul, hybrids may still form but become less viable or rare due to mismatched traits to the environments or flowering times. The two populations are not fully separate species, but they are no longer freely interbreeding either. If environments shift, gene flow between populations might resume and blur their boundaries again.

Situations like this, where two populations live in different environments and only partly stop interbreeding (partial reproductive isolation) are very common in nature. Recognizing these "in-between" cases helps us understand how new species gradually form, and why it is not always clear where one species ends and another species begins.

**How Do Gene Flow and Chromosomal Recombination Affect on Speciation?**
Gene flow and recombination both act to mix genetic material between individuals, and this can make speciation process more complex:

**Gene flow** introduces alleles from one population into another, which tends to homogenize differences and slow down divergence. **Recombination** further breaks up associations between locally adapted alleles, making it harder for advantageous combinations of multiple alleles to persist. However, speciation can still occur when certain evolutionary conditions arise despite these forces. Such conditions may include strong selection against migrants, changes in mating preference that reduce interbreeding, or the emergence of fixed genomic structures, for example, **chromosomal inversions**, that locally suppress recombination.

When gene flow and recombination are reduced in certain parts of the genome, it becomes easier for populations to maintain genetic differences. This allows genetic divergence to build up, even while other parts of the genome remain less diverged between populations. As a result, the early stages of speciation often produce a patchy pattern in the genome, some regions show differentiation, while others do not. These patterns can still remain visible long after the species have fully diverged.

#### Recommended Resource

**Moran et al. (2022) – The genomic consequences of hybridization**
This article provides a genomic perspective on hybridization, showing its widespread role in evolution, adaptation, and speciation across taxa using conceptual figures.

### 2.1 Population Genomics : Demographic History and Evolution

This lecture introduces how we study genetic variation within and between populations using genome-wide data. It explains how we can use genomics to investigate population structure, past demographic events like bottlenecks or migration, and signs of natural selection.
It also discusses problems of current reference-genome based methods and introduces pangenomes as a solution.

#### Vocabulary

1. **Admixture analysis**: Statistical estimation of ancestral population proportions within individuals.
2. **Runs of homozygosity (ROH)**: Long homozygous regions indicating inbreeding or demographic events.
3. **Identity by state (IBS)**: Shared genomic segments regardless of ancestry; used to infer relatedness and demography.
4. **Selective sweep**: A reduction in genetic variation due to rapid fixation of a beneficial allele.
5. **$F_{ST}$ (Fixation index)**: A measure of population differentiation; used to identify divergent regions under selection.
6. **Genotype–Environment Association (GEA)**: Methods linking allele frequency variation to environmental gradients.
7. **Linkage disequilibrium (LD)**: Non-random association of alleles at different loci; affected by recombination, selection, and structure.
8. **Inversions and supergenes**: Structural variants that suppress recombination and preserve co-adapted gene sets.
9. **Reference bias**: Systematic errors arising from aligning reads to a single reference genome, often underestimating variation.
10. **Pangenome**: A genomic framework incorporating core and accessory sequences across individuals, reducing reference bias.

**How Demographic History Affects the Genome**

**Demography**, the size and structure of a population over time has a powerful influence on patterns of genetic variation. Imagine a small island population of birds that survives a typhoon, with only a few individuals left to reproduce. This kind of **bottleneck** reduces genetic diversity, because the gene pool is suddenly restricted to the survivors. If two of these birds happen to carry a rare harmful mutation, future generations may suffer from inbreeding, making it harder to find healthy breeding partners.

In another case, a large fish population may split into two groups when a lake dries up and forms two separate basins. Over time, each group accumulates different mutations and begins to diverge. Conversely, populations that were separated may come back into contact, for instance, when a glacier melts and reconnects rivers, and their genomes begin to mix again, creating mosaic ancestry.

Sometimes, a population expands rapidly, such as when a few individuals colonize a new habitat with plenty of resources. This growth can lead to the accumulation of many new mutations, especially rare ones. These demographic events, bottlenecks, expansions, isolation, and admixture, leave distinct signatures in the genome.

Understanding demography is also crucial in conservation and breeding: a population with low diversity may be more vulnerable to disease or show reduced fertility due to accumulation of deleterious alleles.

##### How Selection Affects the Genome

**Selection** acts on variation in traits, promoting the spread of beneficial variants and removing harmful variants from the population. In **directional selection**, one version of a trait becomes strongly favored: for example, in a desert environment, lighter-colored lizards may avoid predators more easily, so genes for pale coloration spread. **Positive selection** increases the frequency of alleles that provide a fitness advantage, while **negative selection** removes alleles that have harmful effects on survival or reproduction.

In **diversifying selection**, different traits are favored in different settings, for instance, deep-water fish might evolve dark coloration while shallow-water fish evolve white, depending on light conditions. This can lead to genetic divergence even without full geographic separation. **Balancing selection** can preserve diversity, such as when having two different alleles confers resistance to multiple pathogens.

Sometimes, **parallel selection** occurs: the same traits evolve independently in different populations facing similar environments, using the same or different genetic pathways. And of course, **artificial selection**, by humans, can dramatically reshape the genome, as in domestic animals, crop, or in aquaculture, where traits like size, behavior, or reproduction are intentionally selected. Selection produces detectable patterns in the genome.

Selection does not always have strong or easily detectable effects. In the case of **polygenic trait**, which are influenced by many genes, small changes in allele frequencies can accumulate gradually under weak selection. Both natural and artificial selection can influence the genetic composition of populations, depending on which traits increase an individual's reproductive success or are intentionally favored in breeding programs.

#### Find the Answers to These Questions During the Lecture

1. What is population structure, and how do we investigate it?
2. How can we distinguish a signature of selection from one caused by demography, like a bottleneck?
3. Why are some genomic regions more differentiated between populations than others?
4. How does pan genomics help address the limitations of single-reference genome analysis?
5. How do different demographic histories shape genetic diversity in current populations?

### Recommended Resource

**Bourgeois & Warren (2021) – An overview of current population genomics methods for the analysis of whole-genome resequencing data in eukaryotes**

This review introduces key population genomic methods e used for demographic inference and those for detecting selection. You can focus on the broad logic and what kind of phenomenon is detectable in the population genomics and skip over the detailed software comparison tables.

### 2.2 GWAS and eQTLs: Connecting Genotype to Phenotype

This lecture focuses on how genetic variants are linked to trait variation. It explains the idea of heritability (how much of the phenotypic variation can be attributed to genetic variation) and how it can be measured. It introduces QTL mapping and GWAS as methods to find variants associated with traits. It also explains how to handle multiple testing and how to interpret statistical results. Finally, it introduces eQTLs, which link genetic variants to gene expression, helping to identify possible causal effects at the molecular level.

#### Vocabulary

1. **Complex/polygenic traits**: Phenotypes influenced by many genes and environmental factors; contrast with Mendelian (single-gene) traits.
2. **Heritability ($H^2$)**: The proportion of phenotypic variance attributable to genetic variance; may be estimated from pedigree or genomic data.
3. **Quantitative trait locus (QTL)**: A genomic region statistically associated with variation in a quantitative trait, often identified via crossbreeding or association mapping.
4. **Genome-wide association study (GWAS)**: A method to identify common variants statistically associated with traits using genotyping across many individuals.
5. **P-value and multiple testing correction**: Statistical tools to assess significance while accounting for the large number of tests in GWAS (e.g., Bonferroni correction).
6. **Linkage disequilibrium (LD)**: The non-random association of alleles at different loci, often leading to GWAS peaks covering multiple variants.
7. **Causal variant**: A variant that directly affects phenotype or gene expression; often difficult to distinguish from nearby non-causal variants in LD.
8. **Missing heritability**: The discrepancy between heritability estimates from family studies and the fraction explained by GWAS-identified variants.
9. **Expression quantitative trait locus (eQTL)**: A genomic locus associated with gene expression levels; may act in cis (near the gene) or trans (distant or different chromosome).
10. **TPM (Transcripts Per Million)**: A normalized measure of gene expression used in RNA-seq data (see 3.1).

#### Background Concepts

##### Too Many p-values: The Statistical Challenge of Genome-Wide Testing

Genome-wide association studies (GWAS) aim to identify genetic variants associated with specific traits. For example, if we study eye color using data from 10,000 people with blue eyes and 10,000 with brown eyes, we might find that allele A is more commonly observed in blue-eyed individuals. To assess whether this difference is meaningful or just random, we use statistical testing.

**The chi-squared ($\chi^2$) test** compares observed and expected genotype frequencies under the assumption that there is no association (**the null hypothesis**). If the $\chi^2$ value is large and the resulting p-value is very small, we consider **the association statistically significant**. However, when testing many variants (as in GWAS) some p-values will appear small **by chance alone**, even if no true associations exist.

This leads to **the multiple testing problem.** To adjust for it, we apply corrections**. The Bonferroni correction** sets a very strict significance threshold by dividing 0.05 by the number of tests. Although this reduces false positives, it can miss true associations, especially if variants are correlated. An alternative is controlling **the False Discovery Rate (FDR)** using methods like Benjamini–Hochberg, which allows more discoveries while still limiting errors.

For continuous traits, like gene expression in eQTL studies, **linear regression** is used instead of the chi-squared test. Here, we examine whether different genotypes explain differences in expression levels. As in GWAS, multiple testing correction is required due to the large number of comparisons.

#### Find the Answers to These Questions During the Lecture

1. Why is it difficult to map the same trait across different populations?
2. What makes a variant a good candidate for follow-up after GWAS?
3. What are the limitations of GWAS in identifying causal variants?
4. How an eQTL is conducted, and how does it help connect genotype to gene expression?
5. How do cis-eQTLs and trans-eQTLs differ in their genomic context and interpretation?

#### Recommended Resource

**Tam et al. (2019) – Benefits and limitations of genome-wide association studies**
This article presents both the strengths and challenges of GWAS, including its role in trait-associated gene discovery and its limitations in explaining heritability. It is a good conceptual introduction to population-based genetic association studies. Focus on the examples and critique of the GWAS framework.

### 2.3 Genome Editing: Connect Genotype and Mechanism

#### What This Lecture Covers

This lecture explains how genome editing tools including CRISPR/Cas9 are used to test whether genetic variants have real effects.
It introduces knockout, activation, and interference methods, and explains how guide RNAs for these experiments are designed. It also covers how genome-wide CRISPR screens can identify biologically relevant genes and how reporter assays like luciferase tests are used to study regulatory DNA. These methods help connect genetic data to biological function.

#### Vocabulary

1. **Knockout (KO)**: Artificial disruption of gene function.
2. **CRISPR/Cas9**: A genome editing tool that uses guide RNAs to introduce targeted DNA double-strand breaks, leading to gene editing.
3. **CRISPRa / CRISPRi**: Modified CRISPR systems used for activation or repression of gene expression without DNA cleavage (e.g., using dCas9).
4. **Guide RNA (gRNA)**: A synthetic RNA that directs Cas9 to a specific genomic sequence via base pairing.
5. **On-target / Off-target scoring**: Computational assessment of gRNA efficiency and specificity.
6. **Genome-wide CRISPR screens**: High-throughput assays using pooled gRNA libraries to identify functional genes in specific conditions (e.g., GeCKO screens).
7. **Positive / Negative selection screens**: Experimental setups where cells with or without specific edits survive or are depleted under selective pressure.
8. **Reporter assay**: A plasmid-based method to test whether a regulatory region (e.g., promoter variant) alters gene expression.
9. **Luciferase**: An enzyme producing bioluminescence, commonly used as a readout in reporter assays.
10. **Massively parallel reporter assay (MPRA)**: A high-throughput method where thousands of DNA elements are tested for regulatory activity simultaneously.

#### Find the Answers to These Questions During the Lecture

1. How does CRISPR/Cas9 introduce gene knockouts?
2. What factors should be considered when designing a guide RNA (gRNA) for CRISPR experiments?
3. How can we detect whether CRISPR editing was successful?
4. How do luciferase reporter assays help test whether a regulatory variant affects gene expression?
5. What are the advantages of massively parallel reporter assays over single reporter constructs?

#### Background Concepts

##### Genome Editing

**Genome editing** refers to techniques that allow scientists to make precise, targeted changes to the DNA of living cells. These changes can involve inserting, deleting, or replacing specific sequences. Most genome editing tools work by creating a **double-strand break** at a chosen spot in the genome. The cell then attempts to repair the break using its own repair systems, which can be harnessed to introduce desired changes. **CRISPR-Cas9** was adapted from a natural **immune defense system in bacteria**. When bacteria are infected by viruses (called **phages**), they can store pieces of the viral DNA in special genome regions. If the same virus attacks again, the bacteria use that stored sequence to make a **guide RNA**, which helps a **Cas protein** recognize and cut the matching viral DNA. Scientists realized they could reprogram this system: by designing a guide RNA to match any gene of interest, they can direct Cas9 to cut specific parts of the genome in **eukaryotic cells**, allowing for highly flexible genome editing.

##### Genome-Wide Screening

In **genome-wide screening**, researchers often apply a **selective pressure**, such as a drug or environmental stress, to a population of cells that have been genetically edited in many different ways. In a **positive selection screen**, cells with certain genetic edits survive better than others under the treatment, so those edits become **enriched** in the population. This helps identify genes that contribute to resistance or survival. In contrast, a **negative selection screen** looks for edits that cause cells to die or grow poorly under the same conditions, so those edits become **depleted** over time. This helps identify genes that are important for survival or fitness under stress. By comparing the abundance of each edit before and after treatment, researchers can infer which genes play a role in the biological process being studied. These methods are often used with **CRISPR** or **RNAi** libraries that target many genes across the genome.

#### Recommended Resource

**Przybyla & Gilbert (2022) – A new era in functional genomics screens**

A clear and forward-looking review of CRISPR-based screening technologies used to investigate gene function, interactions, and regulatory elements. Useful for students interested in experimental genomics or gene editing.

### 3.1 Current Transcriptomics: Across Space, Time, and Cell Type

#### What This Lecture Covers

This lecture introduces RNA sequencing (RNA-seq) as a way to measure gene expression. It explains how bulk RNA-seq and single-cell RNA-seq differ, and how each method can identify which genes are active in different cells or conditions. It also covers how to analyze RNA-seq data, including normalization and differential expression. The lecture introduces newer methods like spatial transcriptomics, which adds location information, and epitranscriptomics, which looks at chemical modifications of RNA.

#### Vocabulary

1. **RNA-seq**: High-throughput sequencing of cDNA to quantify transcript abundance across genes; the basis of digital gene expression profiling. -> See **0.1**
2. **Library preparation**: Protocol to convert mRNA into cDNA and prepare it for sequencing; may introduce bias (e.g., poly-dT priming).
3. **Read count**: The number of sequencing reads mapping to each gene; used as a proxy for expression level.
4. **Normalization**: Mathematical adjustment to account for sequencing depth or gene length
5. **Differential gene expression**: genes whose expression differs between conditions/groups
6. **Single-cell RNA-seq (scRNA-seq)**: Technique for profiling gene expression at the level of individual cells; enables discovery of rare cell types and trajectories.
7. **Dimensionality reduction**: Methods such as t-SNE or UMAP that allow visualization of transcriptomes in 2D space.
8. **Trajectory inference**: Computational reconstruction of differentiation paths using pseudotime; used in developmental biology.
9. **Spatial transcriptomics**: Combines RNA-seq with spatial localization of transcripts on tissue sections.
10. **Epitranscriptomics**: Study of chemical modifications to RNA (e.g., m6A) that influence transcript stability and translation.

#### Find the Answers to These Questions During the Lecture

1. Why is normalization essential in RNA-seq analysis?
2. How are cell types and developmental paths inferred from scRNA-seq data?
3. What biological insights can be gained from spatial transcriptomics?
4. How do RNA modifications affect gene regulation?
5. What is the role of pseudotime analysis in developmental biology?

#### Background Concepts

**Dimensionality reduction** refers to a set of techniques used to make complex, high-dimensional data easier to explore and interpret, especially visually (Also see **2.1**). For example, when analyzing **transcriptomic** data, each cell or sample can be represented by thousands of gene expression values, meaning each data point exists in a space with thousands of dimensions.

Humans, however, can only visualize in two or three dimensions. Methods like **t-SNE** and **UMAP** project the original data into a two-dimensional space in a way that tries to preserve certain features of the original structure, such as clusters or distances between points. This makes it possible to see, for instance, whether samples form distinct groups or gradients. These methods are especially useful for exploring **heterogeneity** in single-cell data. However, some information is lost in the process, and the shape of the 2D map does not fully reflect the structure of the original high-dimensional data. Therefore, these plots are best used for exploration and hypothesis generation, not as definitive evidence.

#### Recommended Resource

**RNA sequencing: the teenage years** ***- Stark, R., Grzelak, M., & Hadfield, J. (2019)***
This is a comprehensive review of RNA-seq methods, including bulk and single-cell applications, normalization, and analysis workflows.

### 3.2 Genomic Changes in a Body: Somatic Mutation, Cancer, Immune Cells

#### What This Lecture Covers

In this lecture, we will learn about how DNA in an individual's body changes over time. It describes how somatic mutations occur, how they are repaired, and how they can lead to diseases like cancer. It also explains special types of genome changes in immune cells that help create antibody diversity. The lecture discusses how mutation rates differ between tissues and individuals, and how to detect these mutations using sequencing data.

#### Vocabulary

1. **Genome plasticity**: The capacity of a genome to undergo structural or sequence-level changes, either during development or in response to environmental stimuli.
2. **Somatic mutation**: A mutation occurring in a non-germline cell, potentially leading to mosaicism or disease (e.g., cancer).
3. **DNA repair pathways**: Systems that correct DNA damage.
4. **VDJ recombination**: Programmed DNA rearrangement in immune cells that generates antibody diversity.
5. **Chromosomal translocation / duplication**: Rearrangements often seen in cancer; may be spontaneous or mutagen-induced
6. **Insertional mutagenesis**: Disruption of host DNA by exogenous sequences, such as retroviruses or engineered transposons.
7. **Ionizing radiation**: High-energy radiation causing DNA strand breaks and complex lesions.
8. **Reactive oxygen species (ROS)**: Byproducts of cellular metabolism that damage DNA bases and sugar backbone.
9. **Crosslinking agents**: Chemicals that covalently bind DNA strands (e.g., cisplatin), interfering with replication and repair.
10. **Alkylating agents**: Add methyl or ethyl groups to DNA bases, leading to mispairing and mutation.

#### Find the Answers to These Questions During the Lecture

1. What are the main sources of somatic mutations, and how do they differ?
2. How do DNA repair systems function, and what happens when they fail?
3. How do programmed mutational processes differ from accidental mutations?
4. What are mutational signatures, and how are they identified?
5. How do mutation rates vary across tissues, life stages, and species?

#### Background Concepts

**Genome Plasticity**

The genome can be altered by many processes. **Intrinsic genome plasticity** refers to changes that arise from normal cellular activity, including errors in DNA replication, base modifications, or the movement of transposons. These are part of the cell's internal chemistry and are often ongoing. In contrast, **extrinsic genome plasticity** is caused by external environmental factors—such as radiation, toxins, or viral infection, that damage the DNA. Both types of changes can affect how genes work and contribute to genetic diversity.

**Immune-Specific Genome Rearrangements vs General Mutations**
Some genome changes are part of normal, controlled processes. In the immune system, **V(D)J recombination** and **class switching** are mechanisms that intentionally rearrange DNA to produce diverse antibodies. V(D)J recombination happens in developing B and T cells and combines gene segments in different ways to create receptors that recognize many pathogens. Class switching occurs later, changing the type of antibody produced (e.g., from IgM to IgG) without altering its antigen specificity. These processes are examples of **programmed genome plasticity**, they are tightly regulated and essential for adaptive immunity. Unlike random mutations, they occur in specific cell types at specific times, using dedicated enzymes and repair systems.

#### Recommended Resource

**Chen et al. (2025) – Chromosomal instability as a driver of cancer progression**
This article explores how somatic mutation and chromosomal instability contribute to cancer development, metastasis, and treatment resistance.

**Tan et al. (2024) – New insights into antibody structure with implications for self–no self discrimination**
This perspective reveals how regulated genomic rearrangements in immune cells give rise to antibody diversity while minimizing autoimmunity. It exemplifies a functional, highly controlled form of somatic genome plasticity.

### 3.3 Function of the Non-Coding Regions: ATAC-seq, ChIP-seq, and Hi-C:

#### What This Lecture Covers

This lecture focuses on non-coding regions of the genome that control gene expression. It explains how to measure chromatin accessibility using ATAC-seq, how to detect protein–DNA interactions using ChIP-seq, and how to study 3D genome structure with Hi-C. The lecture also introduces the idea of chromatin states and regulatory elements like enhancers and promoters. It shows how these regions are organized and how they affect gene activity in different cell types.

#### Vocabulary

1. **Chromatin accessibility**: Whether genomic DNA is exposed and available for transcription factor binding; measured by ATAC-seq or DNase-seq.
2. **Histone modifications**: Post-translational marks (e.g., H3K4me3, H3K27ac) that indicate active or repressed chromatin states; measured by ChIP-seq.
3. **Enhancer / promoter**: Regulatory DNA elements that activate transcription; promoters are near transcription start sites, enhancers can act at a distance.
4. **Chromatin looping**: Physical interactions between distal elements (e.g., enhancer–promoter) that bring regulatory regions into contact.
5. **Hi-C**: A sequencing method that captures genome-wide chromatin interactions by proximity ligation; used to define 3D structure.
6. **Topologically Associating Domains (TADs)**: Self-interacting chromatin regions with high internal contact frequency; boundaries often conserved.
7. **A/B compartments**: Large-scale active (A) or inactive (B) genomic regions identified through principal component analysis of Hi-C data.
8. **Chromatin states**: Computationally defined regions such as "promoter", "enhancer", or "repressed" based on combined assay signals.
9. **ENCODE project**: A large-scale effort to map regulatory elements across the human and mouse genomes.
10. **SCREEN**: A searchable database of candidate cis-regulatory elements produced by ENCODE.

#### Find the Answers to These Questions During the Lecture

1. How does chromatin structure affect gene expression?
2. How do enhancers activate genes over long genomic distances?
3. What is the difference between chromatin accessibility and histone marking?
4. How are chromatin states defined and annotated?
5. How can comparative regulatory maps be constructed and applied?

#### Background Concepts

In the 3D organization of the genome, chromosomes are not randomly packed inside the nucleus. Instead, they tend to fold into regions with different activity states. One way to detect this organization is by using **Hi-C**. **Hi-C** is a technique used to measure how often different parts of the genome are physically close to each other in the 3D space of the nucleus. The result is a large matrix, where each cell indicates how frequently two genomic regions interact. For example, if two regions are often found near each other in the nucleus, their interaction frequency will be high.

When this matrix is visualized, it often reveals patterns, some regions interact mostly with themselves, forming squares along the diagonal, while others show broader interaction patterns.

To detect large-scale organization, researchers apply **principal component analysis (PCA)** to this matrix (see 2.1). PCA helps identify broad trends in the interaction data, such as whether a region tends to interact more with "active" or "inactive" areas. The first principal component often separates the genome into **A compartments** (active, gene-rich) and **B compartments** (inactive, gene-poor), revealing a functional organization of the genome based on how it folds in 3D space.

#### Recommended Resource

**Oudelaar & Higgs (2021) – The relationship between genome structure and function**
This review discusses how the 3D structure of the genome influences gene, bridging chromatin architecture and gene expression regulation.

### 4.0 Final Project – Research Proposal and Presentation

This module gives you the opportunity to integrate concepts, data types, and methods introduced throughout the course into a hypothetical research proposal. Your task is not only to propose a biologically interesting question, but also to think as a scientist: to formulate testable hypotheses, consider risks and limitations, and design a study that reflects ambition and feasibility.

#### Key questions to guide your thinking include:

1. What is the **state of the art** in the field related to your question?
2. What is your **hypothesis** and what are its implications?
3. What kind of **data and analyses** would provide the most insight?
4. What are the **risks** to your project's success, and how might you mitigate them?
5. What is the **potential impact** of your findings, within or beyond biology?

#### Vocabulary

1. **Research question**: The broader question that motivates the project, often open-ended.
2. **State of the art**: The current knowledge frontier in a particular field. Your project should define its position relative to this.
3. **Gap in knowledge**: An area where current scientific understanding is incomplete or missing, justifying the need for further research
4. **Hypothesis**: A clear, testable statement that links cause and effect or mechanism to pattern.
5. **Objective**: A concrete, achievable target that addresses the hypothesis.
6. **Methodology**: The rationale for choosing specific techniques, tools, or data types. Should be feasible and well justified.
7. **Risk and mitigation**: Identification of factors that may hinder the project, with strategies to reduce or manage them.
8. **Feasibility**: Whether the question can be addressed given the data, time, and expertise available.
9. **Novelty**: What is original in your question, approach, or expected result. Not just "new" but **unexpected and informative**.
10. **Impact**: The potential for your findings to shape understanding, methods, or applications—either within genomics or in broader science.

#### Recommended Resource

**Weidmann et al. (2023) – How to write a successful grant application**
This commentary provides a structured overview of key elements in writing grant proposals, from conceptualizing the research aim to articulating hypotheses and selecting appropriate methods. For our purposes, you can focus on how the paper emphasizes clarity in research questions and methodological logic, while disregarding sections on budgeting, funding schemes, or timeline planning, which are not relevant to this assignment.
